# The lncRNA *MARS* modulates the epigenetic reprogramming of the marneral cluster in response to ABA

**DOI:** 10.1101/2020.08.10.236562

**Authors:** Thomas Roulé, Federico Ariel, Caroline Hartmann, Nosheen Hussain, Moussa Benhamed, Jose Gutierrez-Marcos, Martin Crespi, Thomas Blein

## Abstract

Clustered organization of biosynthetic non-homologous genes is emerging as a characteristic feature of plant genomes. The co-regulation of clustered genes seems to largely depend on epigenetic reprogramming and three-dimensional chromatin conformation. Here we identified the long noncoding RNA (lncRNA) *MARneral Silencing* (*MARS*), localized inside the Arabidopsis marneral cluster, which controls the local epigenetic activation of its surrounding region in response to ABA. *MARS* modulates the POLYCOMB REPRESSIVE COMPLEX 1 (PRC1) component LIKE-HETEROCHROMATIN PROTEIN 1 (LHP1) binding throughout the cluster in a dose-dependent manner, determining H3K27me3 deposition and chromatin condensation. In response to ABA, *MARS* decoys LHP1 away from the cluster and promotes the formation of a chromatin loop bringing together the *MARNERAL SYNTHASE 1* (*MRN1*) locus and a distal ABA-responsive enhancer. The enrichment of co-regulated lncRNAs in clustered metabolic genes in Arabidopsis suggests that the acquisition of novel noncoding transcriptional units may constitute an additional regulatory layer driving the evolution of biosynthetic pathways.

## INTRODUCTION

In eukaryotes, functionally related genes are usually scattered across the genome. However, a growing number of operon-like clustered organization of non-homologous genes participating in common metabolic pathways point at an emerging feature of animal, fungi and plant genomes (1).

In plants, synthesis of numerous secondary metabolic compounds is important for the dynamic interaction with their environment, affecting their life and survival (2). Terpenoids are bioactive molecules of diverse chemical structure (3). In *Arabidopsis thaliana*, the biosynthesis of four triterpenes, namely thalianol (4), tirucalla-7,24-dien-3b-ol (5), arabidiol (6) and marneral (7), is governed by enzymes encoded by genes organized in clusters (1). The thalianol and marneral related genes are located in the smallest metabolic clusters identified in plants to date, each being less than 40kb in size (1). Both compounds are derived from 2,3-oxidosqualene and the corresponding gene clusters contain the oxidosqualene cyclases (OSCs), thalianol synthase (THAS) and marneral synthase (MRN1), respectively. The marneral cluster includes two additional protein-coding genes, *CYP705A12* and *CYP71A16*, participating in marneral oxidation (7).

Growing evidence indicates that the co-regulation of clustered genes relies on epigenetic mechanisms. It has been shown that the deposition of the histone variant H2A.Z positively correlates with transcriptionally active clusters. Accordingly, nucleosome stability precluding gene expression is dependent on ARP6, a component of the SWR1 chromatin remodeling complex required for the deposition of H2A.Z into nucleosomes (8). Additionally, it was shown that the thalianol and marneral clusters exhibit increased expression in the Polycomb mutant *curly leaf* (*clf*) with compromised H3K27me3 deposition, and reduced expression in the trithorax-group protein mutant *pickle* (*pkl*), a positive regulator that counteracts H3K27me3 silencing (9). Strikingly, it has been recently shown that biosynthetic gene clusters are embedded in local hot spots of three-dimensional (3D) contacts that segregate cluster regions from the surrounding chromosome environment in a tissue-dependent manner. Notably, H3K27me3 appeared as a central feature of the 3D domains at silenced clusters (10).

Long noncoding RNAs (lncRNAs) have emerged as important regulators of eukaryotic gene expression at different levels (11). In plants, several lncRNAs have been shown to interact with the Polycomb Repressive Complex 1 and 2 components LIKE HETEROCHROMATIN PROTEIN 1 (LHP1) and CLF, respectively, which are related to H3K27me3 distribution (12, 13). Furthermore, it has been proposed that lncRNAs can modulate the transcriptional activity of neighboring genes by shaping local 3D chromatin conformation (14–16). Here we show that the marneral cluster in Arabidopsis includes three noncoding transcriptional units. Among them, the lncRNA *MARS* influences the expression of marneral cluster genes in response to ABA through modification of the epigenetic landscape. *MARS* deregulation affects H3K27me3 distribution, LHP1 deposition and chromatin condensation throughout the cluster. Furthermore, an ABA responsive chromatin loop dynamically regulates *MRN1* transcriptional activation by bringing together the *MRN1* proximal promoter and an enhancer element enriched in ABA-related transcription factors (TF) binding sites. *MARS*-mediated control of the marneral cluster affects seed germination in response to ABA. The general co-regulation of genes located within lncRNA-containing clusters in Arabidopsis points to noncoding transcription as an important feature in coordinated transcriptional activity of clustered loci.

## MATERIAL AND METHODS

### Lines selection and generation

All plants used in this study are in Columbia-0 background. RNAi-*MARS* were obtained using the pFRN binary vector (17) bearing 250bp of the first exon of *MARS* gene (see primers in **Supplementary Table 1**), previously sub-cloned into the pENTR/D-TOPO vector. Arabidopsis plants were transformed using *Agrobacterium tumefaciens* Agl-0 (19). The T-DNA inserted line *SALK_133089* was ordered to NASC (N633089). Homozygous mutants were identified by PCR (see primers in **Supplementary Table 1**).

Seeds of *rnm1* (2), *35S:MRN1* (7), and *mro1-2* (*cyp71a16*, (7)) mutants were kindly provided by Dr. Ben Field (BIAM, CEA Cadarache, France) and Pr. Suh (Chonnam National University, Department of Bioenergy Science and Technology, Korea), respectively.

### Growth conditions and phenotypic analyses

Seeds were sown in plates vertically placed in a growing chamber in long day conditions (16 h in light 150uE; 8 h in dark; 21°C) for all the experiments. Plants were grown on solid half-strength MS medium (MS/2) supplemented with 0.7% sucrose, and without sucrose for the germination assay. For nitrate starvation assay, KNO_3_ and Ca(NO_3_)_2_ were replaced from MS/2 by a corresponding amount of KCl and CaCl_2_ respectively, 2.25 mM NH_4_HCO_3_ was added for nitrate-containing medium. For the phosphate starvation assay, growth medium contained 0.15 mM MgSO_4_, 2.1 mM NH_4_NO_3_, 1.9 mM KNO_3_, 0.34 mM CaCl_2_, 0.5 μM KI, 10 μM FeCl_2_, 10 μM H_3_BO_3_, 10 μM MnSO_4_, 3 μM ZnSO_4_, 0.1 μM CuSO_4_, 0.1 μM CoCl_2_, 0.1 μM Na_2_MoO_4_, 0.5 g.L^-1^ sucrose supplemented with 500µM Pi for Pi containing medium versus 10µM for Pi free medium. All media were supplemented with 0.8g/L agar (Sigma-Aldrich, A1296 #BCBL6182V) and buffered at pH 5.6 with 3.4mM 2-(N-morpholino) ethane sulfonic acid. For the treatment with water, exogenous ABA or auxin, seedlings were sprayed with 10µM to 100µM ABA and 10µM 1-Naphthaleneacetic acid (NAA), respectively. For heat stress, plates were transferred to a growth chamber at 37°C under the same lightning conditions. For nitrate and phosphate starvation assays, seedlings were transferred at day 12 after sowing (DAS) from respectively nitrate and phosphate containing medium to nitrate and phosphate free medium. For root phenotype characterization, seedlings were sown in control media and transferred at day 6 in control medium or medium containing 2µM ABA, 200mM mannitol or 100mM NaCl, respectively. After 3 days of growth, the root length was measured using RootNav software (20) from images taken with a flat scanner. Finally, for seed germination assay, 0.5µM ABA was supplemented or not to the medium. Germination rate was evaluated twice a day. Seeds were considered germinated when the seed coat was perforated by elongating radicle. For all the experiments, samples were taken from 12 DAS starting two hours after light illumination, at different time-points, after cross-linking or not, depending on the experiment.

### RT-qPCR

Total RNA was extracted from whole seedlings using TRI Reagent (Sigma-Aldrich) and treated with DNase (Fermentas) as indicated by the manufacturers. Reverse transcription was performed using 1µg total RNA and the Maxima Reverse Transcriptase (Thermo Scientific). qPCR was performed on a Light Cycler 480 with SYBR Green master I (Roche) in standard protocol (40 cycles, 60°C annealing). Primers used in this study are listed in **Supplementary Table 1**. Data were analyzed using the ΔΔCt method using *PROTEIN PHOSPHATASE 2A SUBUNIT A3* (*AT1G13320*) for gene normalization (21) and time 0 for time-course experiments.

### Chromatin Immunoprecipitation (ChIP)

ChIP was performed using anti-IgG (Millipore,Cat#12-370), anti-H3K27me3 (Millipore, Cat#07-449) and anti-LHP1 (Covalab, Pab0923-P), as previously described (14), starting from two grams of seedlings crosslinked in 1% (v/v) formaldehyde. Chromatin was sonicated in a water bath Bioruptor Plus (Diagenode; 60 cycles of 30s ON and 30s OFF pulses at high intensity). ChIP was performed in an SX-8G IP-Star Compact Automated System (Diagenode). Antibody-coated Protein A Dynabeads (Invitrogen) were incubated 12 hours at 4 °C with the samples. Immunoprecipitated DNA was recovered using Phenol:Chloroform:Isoamilic Acid (25:24:1, Sigma) followed by ethanol precipitation and quantified by qPCR. For input samples, non-immunoprecipitated sonicated chromatin was processed in parallel.

*In-vitro* transcribed *MARS and GFP* RNA were obtained from a PCR product amplified from wild-type genomic cDNA and pB7FWG2 plasmid harbouring the *GFP* gene, respectively, using the T7 promoter included in the forward primer used for amplification (**Supplementary Table 1**). PCR products were purified by agarose electrophoresis and NucleoSpin kit (Macherey-Nagel). 1µg of purified DNA was used for *in-vitro* transcription following the manufacturer instructions (HiScribe T7 High Yield RNA Synthesis Kit, NEB). Purified non-crosslinked chromatin obtained from five grams of *MARS* RNAi line 1 seedlings were resuspended in 1 mL of nuclei lysis buffer and split into five tubes. An increasing amount of *MARS* RNA was added to each tube from 0 to 10 µg RNA and incubated under soft rotation during 3 h at 4 °C. Chromatin samples were then cross-linked using 1% (v/v) of formaldehyde for five minutes. Sonication and the following ChIP steps were performed as described above.

### Formaldehyde-Assisted Isolation of Regulatory Elements (FAIRE)

FAIRE was performed as described by (22). After chromatin purification following the ChIP protocol, only 50 µl from the 500 µl of purified chromatin were used (diluted to 500 µl in 10 mM Tris-HCl pH 8). Quantification was performed by qPCR using the same set of primers as for ChIP.

### Immunoprecipitation of methylated DNA (meDIP)

MeDIP was performed as described by (23). For genomic DNA purification, 100mg of non-cross-linked seedlings were incubated 30min at 65°C in 600uL of cetyltrimethylammonium bromide (CTAB) buffer (2% CTAB, 1.4M NaCl, 100mM Tris pH8, 20mM EDTA and 0.2% B-mercaptoethanol). A Chloroform:Isoamyl Alcohol (24:1) wash was performed prior to precipitation with isopropanol. After RNAse A treatment, 1µg of pure DNA was sonicated in a water bath Bioruptor Plus (Diagenode; 10 cycles of 30s ON and 30s OFF pulses at low intensity). The IP of the methylated DNA was performed overnight at 4°C using Protein A Dynabeads coated with anti-5mC (Diagenode, C15200081) or anti-IgG (Diagenode, C15400001). Immunoprecipitated DNA was recovered using Phenol:Chloroform:Isoamyl Alcohol (25:24:1, Sigma) followed by ethanol precipitation and quantified by qPCR. For input samples, non-immunoprecipitated sonicated chromatin was processed in parallel.

### Nuclear purification

Non-cross-linked seedlings were used to assess the sub-cellular localization of RNAs. To obtain the nuclear fraction, chromatin was purified as for ChIP and resuspended, after the sucrose gradient, into 1mL of TRI Reagent (Sigma-Aldrich). For the total fraction, 200 µL of cell suspension from the first step of the ChIP protocol, were treated with 800 µL of TRI Reagent to follow with the RNA extraction. RNA samples were treated with DNase, and RT was performed using random hexamers prior to qPCR analysis.

### RNA immunoprecipitation (RIP)

For RIP, the *lhp1* mutants complemented with the *ProLHP1:LHP1:GFP* (24) were treated for 4h with ABA. After crosslinking and chromatin extraction as for ChIP, ten percent of resuspended chromatin was kept at −20 °C as the input. Chromatin was sonicated in a water bath Bioruptor Plus (Diagenode; 5 cycles of 30 s ON and 30 s OFF pulses at high intensity). Anti-LHP1 RIP was performed using the anti-GFP antibody (Abcam ab290), as previously described (14). The enrichment was determined as the percentage of cDNA detected after IP taking the input value as 100%.

### Chromosome conformation capture (3C)

3C was performed as previously described (25). Briefly, chromatin was extracted from two grams of cross-linked seedlings as for ChIP. Overnight digestion at 37 °C was performed using 400U of Hind III enzyme (NEB). Digested DNA was ligated during 5 h incubation at 16 °C with 100 U of T4 DNA ligase (NEB). DNA was recovered after reverse crosslinking and Proteinase K treatment (Invitrogen) by Phenol:Chloroform:Isoamyl Acid (25:24:1; Sigma) extraction and ethanol precipitation. Interaction frequency was determined by qPCR using a DNA region uncut by Hind III to normalize the amount of DNA across samples.

### Transcriptional activation assay in tobacco leaves

The *GUS* reporter system for validating the activity of the putative enhancer element was adapted from (26). Different DNA fragments were cloned in the GreenGate system (18) fused to a minimal 35S promoter element from CAMV (synthesized by Eurofins Genomics). The sub-unit B3 from 35S promoter element from CAMV (27) was synthesized and used as a positive control. All primers used for cloning are indicated in **Supplementary Table 1**.

*A. tumefaciens*-mediated transient transformation was performed on 5-week-old tobacco plants using a needle-less syringe. Together with enhancer constructs, another vector containing mCherry driven by 35S promoter was co-transfected to control the transformation efficiency. Two leaf discs were collected near the infiltration site. One, to determine the transfection efficiency by mCherry fluorescence observation under epifluorescent microscope. The second was used for GUS staining, as previously described (28). In addition, a single leaf was also stained with the different constructs to compare their relative activity. Samples were incubated 4 h in the dark at 37 °C before observation.

### Identification of lncRNA loci in Arabidopsis gene clusters

The genes of co-expressed clusters were retrieved from (9). The boundaries of the gene clusters were extracted using Araport11 annotations. The boundaries of the metabolic clusters were extracted from the plantiSMASH predicted clusters on Arabidopsis (29). Using Araport11 GFF, the lncRNAs (genes with a locus type annotated as “long_noncoding_rna”, “novel_transcribed_region” or “other_rna”) present within the boundaries of the cluster were retrieved.

### Gene expression correlation analyses

To compute the correlation of expression in different organ of Arabidopsis we used the 113 RNA□seq datasets previously considered for the Araport11 annotations (30). These datasets were generated from untreated or mock□treated wild□type Col□0 plants. After removing the adaptors with Trim Galore with default parameters, the reads were mapped on TAIR10 with STAR v2.7.2a (31) and the parameters ‘╌alignIntronMin 20 ╌alignIntronMax 3000’. Gene expression was then quantified with featureCounts v2.0.0 (32) with the parameters “-B -C -p -s 0” using the GFF of Araport11. Raw counts were then normalized by median of ratios using the DESeq2 R package (33).

For the correlation of expression inside the marneral cluster, the transcript levels of the genes included in the cluster and 25kb around it (four genes upstream and two downstream) were considered for the correlation analysis. Pearson’s correlations for each pair of genes were computed after log 2 transformation of the normalized counts. The correlation value and associated p-value were plotted with the corrplot R package (34).

Inside each co-expressed and metabolic clusters of genes, Pearson’s correlation was computed between every possible pair of lncRNA and coding gene as well as for the genes inside the marneral cluster. The maximum correlation value was kept as an indication of lncRNAs correlation with the genes of the cluster.

### Quantification and statistical analyses

For all the experiments, at least two independent biological samples were considered. For RT-qPCR, each sample was prepared from a pool of 5 to 10 individual seedlings. For biochemistry assays (ChIP, FAIRE, nuclear purification, RIP and 3C) two to five grams of seedlings were prepared for each independent biological sample. For validation of enhancer function, the four leaf discs were taken from four independent tobacco plants. An additional replicate was performed on three independent tobacco plants upon the agroinfiltration of all the different constructs on the same leaf. The tests used for statistical analyses are indicated in the respective figure legends. Statistical tests and associated plots have been generated using R software (v3.6.3(R Core, 2004)) with the help of the tidyverse package (35).

## RESULTS

### The marneral gene cluster contains three noncoding transcriptional units

The small marneral cluster includes three genes: marneral synthase (*MRN1*), *CYP705A12* and *CYP71A16* that are two P450 cytochrome-encoding genes (**Figure 1A**), all participating in the biosynthesis and metabolism of the triterpene marneral (7).

**Figure 1.**
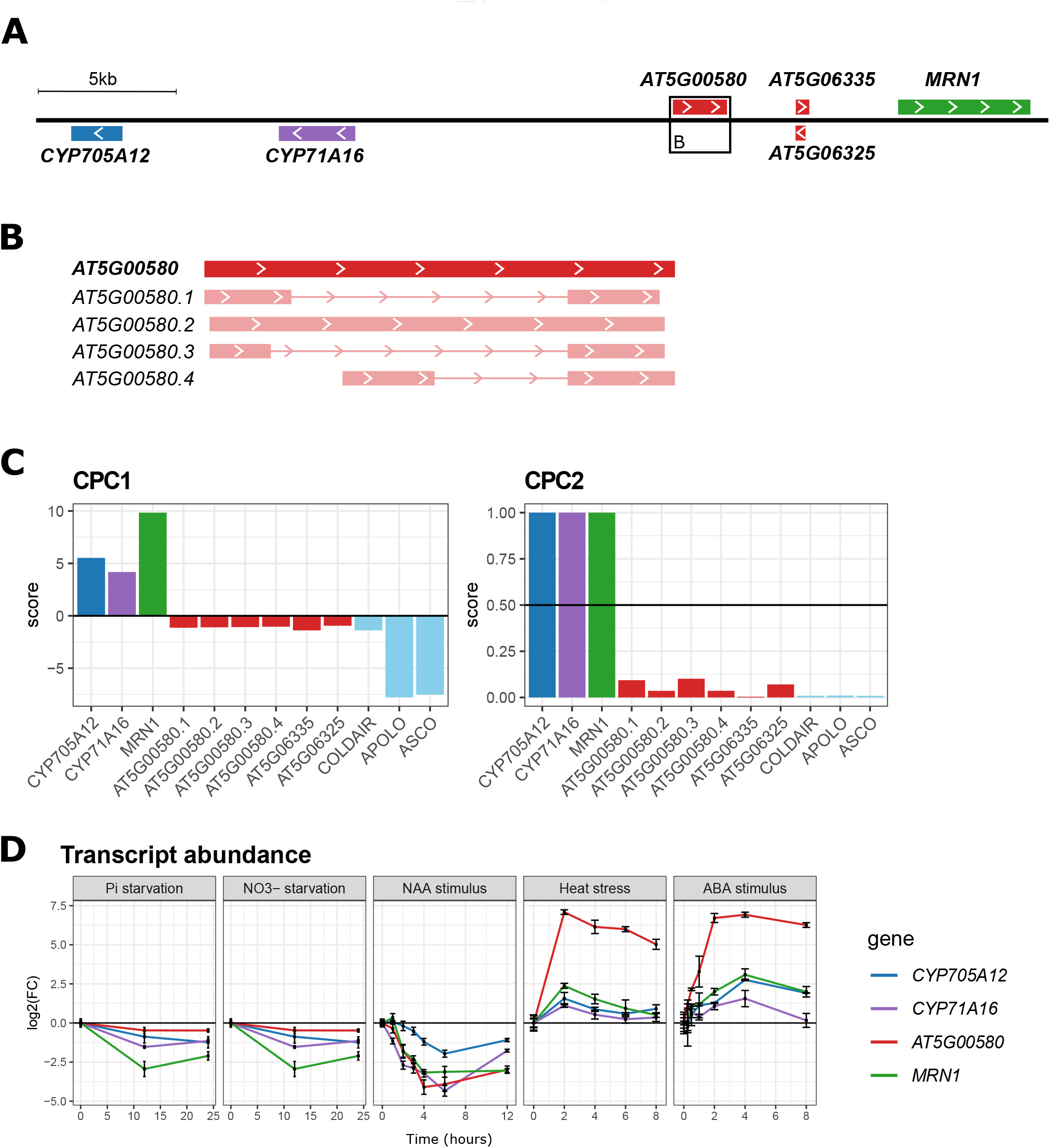
*AT5G00580* is a lncRNA transcribed from the marneral cluster locus and its expression correlates with its neighboring genes. (A) Schematic illustration of the marneral cluster. Genes are indicated with a plain rectangle and white arrows indicate the sense of transcription. The square indicates the region displayed in (B). (B) Schematic illustration of the different isoforms of *AT5G00580* transcripts. First line corresponds to the *AT5G00580* genomic region whereas the other lines present the various isoforms. For each isoform, exons are indicated with rectangle and introns with solid lines. (C) Coding potential of the transcripts located in the marneral cluster genomic region. Scores were determined using CPC1 (left) and CPC2 (right) algorithms (36, 37). For each, the threshold between coding and noncoding genes is displayed with a horizontal solid black line. Coding genes are situated above the threshold, whereas non coding genes are situated under. *COLDAIR, APOLO* and *ASCO* are used as positive controls of noncoding transcripts. (D) Dynamic transcriptional levels of co-regulated genes of the marneral cluster in response to phosphate and nitrate starvation, heat stress, and exogenous ABA and auxin. Gene expression data are shown as the mean ± standard error (n = 3) of the log2 fold change compared to time 0h.

The advent of novel sequencing technologies has allowed the identification of an increasing number of lncRNAs throughout the *Arabidopsis* genome. According to the latest annotation (Araport 11 (30)), three additional transcriptional units are located within the marneral cluster, between the *CYP71A16* and the *MRN1* loci. The *AT5G00580* and the pair of antisense genes *AT5G06325* and *AT5G06335* are located upstream of the *MRN1* gene at 6kpb and 3kbp, respectively (**Figure 1A**). The 1,941bp-long *AT5G00580* locus generates four transcript isoforms ranging from 636 nt to 1,877 nt in length (**Figure 1B**). In contrast, each of the antisense genes *AT5G06325* and *AT5G06335* are transcribed into only one RNA molecule of 509 nt and 367 nt, respectively (**Figure 1A**). All these transcripts were classified as lncRNAs when using two coding prediction tools, CPC (36) and CPC2 (37) because of their low coding potential and their length (over 200 nt), similarly to previously characterized lncRNAs (*COLDAIR* (38); *APOLO* (14); and *ASCO* (39)) (**Figure 1C**).

According to available transcriptomic datasets (Araport11), *AT5G00580* transcriptional accumulation positively correlates with that of marneral genes, whereas *AT5G06325* and *AT5G06335* RNAs do not (**Supplementary Figure S1**). Notably, our analysis of the transcriptional dynamics of the noncoding gene *AT5G00580* and the marneral cluster protein-coding genes revealed a correlated expression in response to phosphate and nitrate starvation, heat stress, as well as to exogenous auxin and ABA (**Figure 1D**). Interestingly, the *AT5G00580* lncRNA exhibited the strongest transcriptional induction in response to heat stress and exogenous ABA, in comparison with *MRN1* and the two *CYP* genes (**Figure 1D**). Altogether, our observations uncovered that the marneral cluster includes three noncoding transcriptional units, one of which is actively transcribed and co-regulated with its neighboring protein-coding genes.

### The lncRNA *MARS* shapes the transcriptional response of the marneral gene cluster to ABA

It has been shown that lncRNAs can regulate the expression of their neighboring genes through epigenetic mechanisms (40). Thus, we wondered if the lncRNA derived from the *AT5G00580* locus may regulate the transcriptional activity of the protein-coding genes included in the marneral cluster. To this end, we modified the lncRNA expression without affecting the cluster DNA region by using an RNAi construct targeting *AT5G00580* transcripts, and isolated three independent lines. The RNAi lines impaired the transcriptional accumulation of *AT5G00580* without affecting significantly the rest of the cluster (**Figure 2A**). Notably, RNAi-mediated silencing did not trigger DNA methylation over the AT5G00580 locus (**Figure S2**). Strikingly, the response of the three protein-coding genes of the marneral cluster to exogenous ABA was significantly deregulated in the RNAi lines (**Figure 2B and *S3A***) compared to mock treatment (**Figure S5**), in contrast to the expression of two *AT5G00580*-unrelated ABA marker genes taken as a control of ABA treatment (**Figure S3B *and* S3C**). Therefore, we named *the AT5G00580*-derived noncoding transcript *MARneral Silencing* (*MARS*) lncRNA. Transcriptional levels of *MRN1* and the two *CYP* genes increased earlier in RNAi-*MARS* seedlings (15 min) than in the wild-type (Col-0, 30 min) (**Figure 2B bottom panel**). In addition, the transcriptional accumulation of these genes later reached two-fold higher levels in the RNAi-*MARS* lines compared to Col-0 (**Figure 2B top panel and S3A**). The same behavior was observed using a higher concentration of ABA (**Figure S4**). Notably, none of the marneral cluster genes exhibit any transcriptional oscillation during the day (41–43), indicating that the transcriptional modulation of *MARS* and the marneral gene cluster linked to an ABA-mediated pathway.

**Figure 2.**
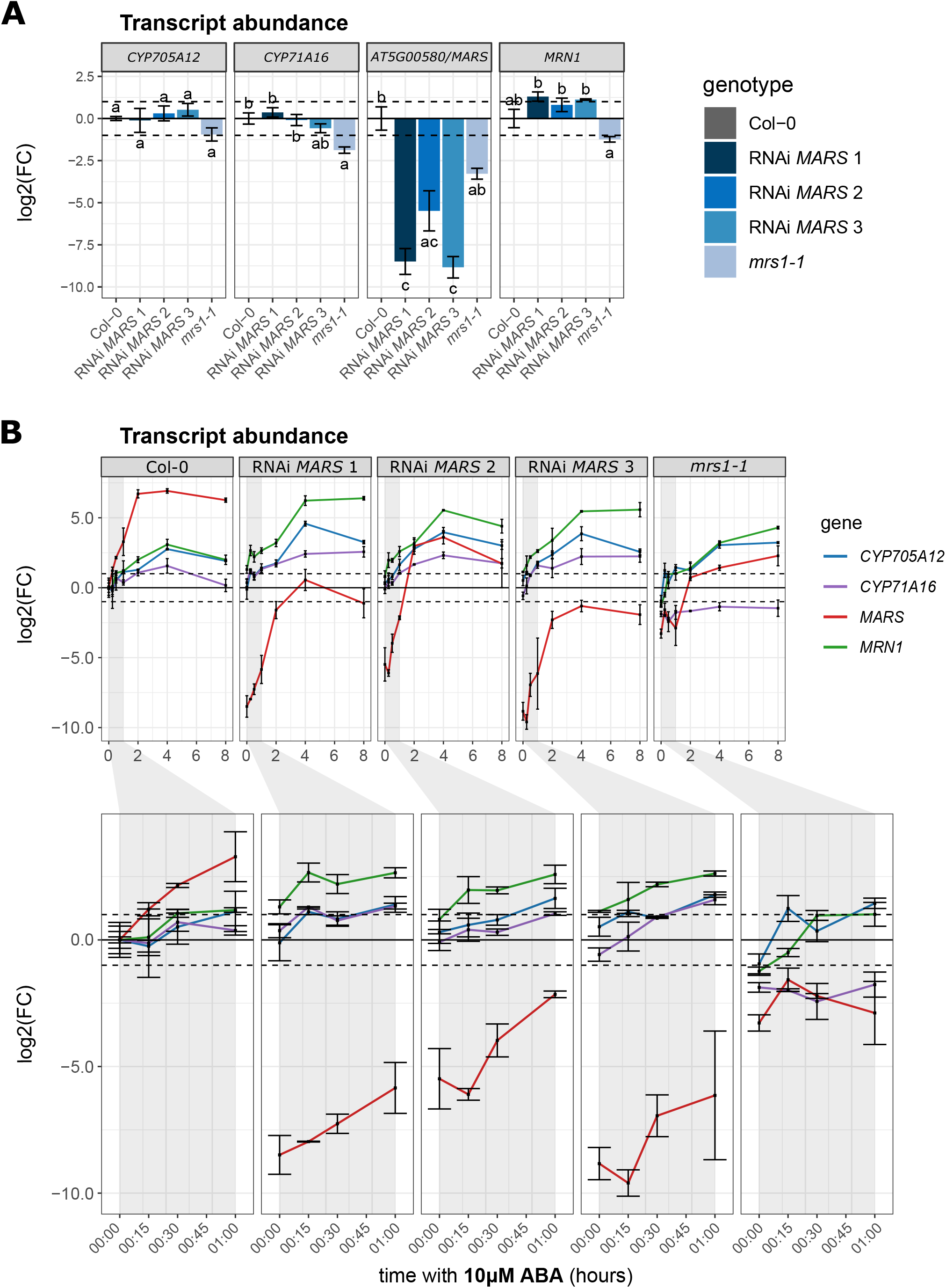
*MARS* transcriptional activity modulates the response to ABA of the marneral cluster. (A) Transcript abundance of the marneral cluster genes in control conditions in RNAi lines targeting *AT5G00580*/*MARS* and *mrs1-1* (*SALK_133089)*. Transcriptional abundance is shown as the mean ± standard error (n = 3) of the log2 fold change compared to the Col-0 genotype. Letters indicate a statistical group determined by one-way analysis of variance (ANOVA) followed by Tukey’s post-hoc test. For each gene, each letter indicates statistical difference between genotypes (p ≤ 0.05). (B) Transcript levels of the genes of the marneral cluster in response to ABA treatment in RNAi lines targeting *AT5G00580*/*MARS* and *mrs1-1* (*SALK_133089*). Gene expression data are shown as the mean ± standard error (n = 3) of the log2 fold change compared to the Col-0 genotype at time 0h.

To further support our observations, we isolated a transgene insertional mutant (*SALK_133089)* located 200 bp upstream the transcription start site (TSS) of *MARS* gene that we named *mrs1-1*. We found that *mrs1-1* partially impairs the transcriptional accumulation of *MARS* and protein-coding genes in the marneral cluster, mainly *CYP71A16* (Figure 2A). In agreement with the RNAi-*MARS* lines, compared to wild-type plants *MRN1* and *CYP705A12* genes responded earlier to ABA and reached higher levels in mrs1-1 (Figure 2B). In contrast, *CYP71A16*, whose promoter region may be locally affected by the T-DNA insertion, was no longer responsive to ABA. In addition, we characterized a transgene insertional mutant in *CYP71A16 gene* (*mro1-2;* (7)), which did not influence the expression of the other marneral cluster genes (**Figure S6A**,**B**) or known ABA-responsive genes (**Figure S6C**,**D**), suggesting that the transcriptional misregulation observed in *mrs1-1* mutant could be caused by the down-regulation of this lncRNA. In agreement, the deregulation of *MRN1* in over-expression and in knock-out mrn1 mutant did not affect the expression of the *CYP* genes of the marneral cluster, nor ABA-responsive genes (**Figure *S7***), but resulted in a decrease in *MARS* transcripts abundance under control condition (**Figure S7A**,**B**). Collectively, our results indicate that the noncoding transcriptional activity of *MARS*, represses the dynamic expression of the marneral cluster genes, mainly *MRN1*, in response to ABA.

### *MARS* affects seed germination and root growth under osmotic stress

The phytohormone ABA has been implicated in the perception and transduction of environmental signals participating in a wide range of growth and developmental events such as seed development, germination and root growth response to environmental stimuli (44).

Considering that the marneral cluster exhibited a strong *MARS*-dependent response to ABA, we wondered what was the physiological impact of *MARS* deregulation during seed germination. We assessed seed germination in Col-0 and *MARS* down-regulated lines with or without exogenous ABA. Notably, *MARS* silencing resulted in a delayed germination compared to wild type seeds, both in response to ABA and in control conditions as revealed by an increase in T50 (time for 50% of germination; **Figure S8A**,**B**,**C**,**D**). Accordingly, *35S:MRN1* and *mro1-2* seeds also exhibit a delayed germination phenotype regardless of the treatment with ABA, whereas the germination speed rate of *mrn1* was only impaired in response to ABA (**Figure S8A**,**B**). The physiological behavior of the cluster-related mutants suggests that the misregulation of marneral genes in the *MARS* down-regulated lines could be linked to an increased sensitivity to ABA during germination (**Figure 2B**).

Considering the influence of the marneral cluster in the regulation of seed germination, we decided to assess root growth response to ABA and ABA-related environmental stimuli such as osmotic stress and salt. Except for *mrs1-1* which showed a significantly reduced root growth, all the other knock-down and mutant lines tested were not affected in root growth under normal growth conditions. However, all the genotypes presented a tendency to reduce root growth in response to ABA or hyperosmotic salt stress (**Figure S8E**). Notably, in response to osmotic stress mediated by mannitol, MARS knock-down lines and the lines with modified *MRN1* expression exhibited a weaker impact on lateral root development with increased lateral root density and length (**Figure S8E;** Lateral root length and Lateral root density). The similar behavior between *MRN1* deregulated lines and RNAi-*MARS* lines suggest that the decreased sensitivity to osmotic stress observed in the RNAi-*MARS* lines could be linked to *MRN1* misregulation. Collectively, our results indicate that *MARS* can modulate various ABA-related physiological responses, through the regulation of *MRN1* expression.

### *MARS* controls the epigenetic status of the marneral locus

It has been shown that gene clusters in plants are tightly regulated by epigenetic modifications, including the repressive mark H3K27me3 (9). ChIP-Seq datasets (45) reveals that the marneral cluster region is highly enriched in H3K27me3 in shoots and overlaps with the deposition of LHP1 (**Figure S9)**. ATAC-Seq data (46) also revealed that the marneral cluster exhibits a high chromatin condensation in shoots (**Figure S9**). These data suggest that the marneral cluster is in a epigenetically silent state in aerial organs, thus correlating with its low expression level in leaves (9).

We wondered if the transcriptional activation of the marneral cluster in response to exogenous ABA was associated with a dynamic epigenetic reprogramming. We first assessed H3K27me3 deposition across the marneral cluster, including the gene body of *MRN1, MARS* and the two *CYP* loci (**Figure 3A and S10**). Interestingly, exogenous ABA triggered a strong reduction of H3K27me3 deposition throughout the marneral cluster (**Figure 3A and S10**). Markedly, H3K27me3 basal levels were also significantly lower in RNAi-*MARS* seedlings. Remarkably, H3K27me3 deposition was even lower across the body of all genes of the cluster in response to ABA in the RNAi-*MARS* lines when compared with Col-0, in agreement with the stronger induction by ABA of this subset of genes upon *MARS* silencing (**Figure 3A, S10 and Figure 2B**). Furthermore, we assessed the deposition of LHP1 on different regions of the marneral cluster and found that LHP1 was enriched at the MRN1 promoter and more weakly across *MARS* gene body and the intergenic region between *CYP71A16* and *MARS* (**Figure 3B and S11**). Remarkably, LHP1 recognition was strongly impaired in response to ABA as well as in RNAi-*MARS* seedlings (**Figure 3B and S11**). Therefore, our results indicate that ABA triggers an epigenetic reprogramming of the marneral cluster, likely in a process involving the lncRNA *MARS*.

**Figure 3.**
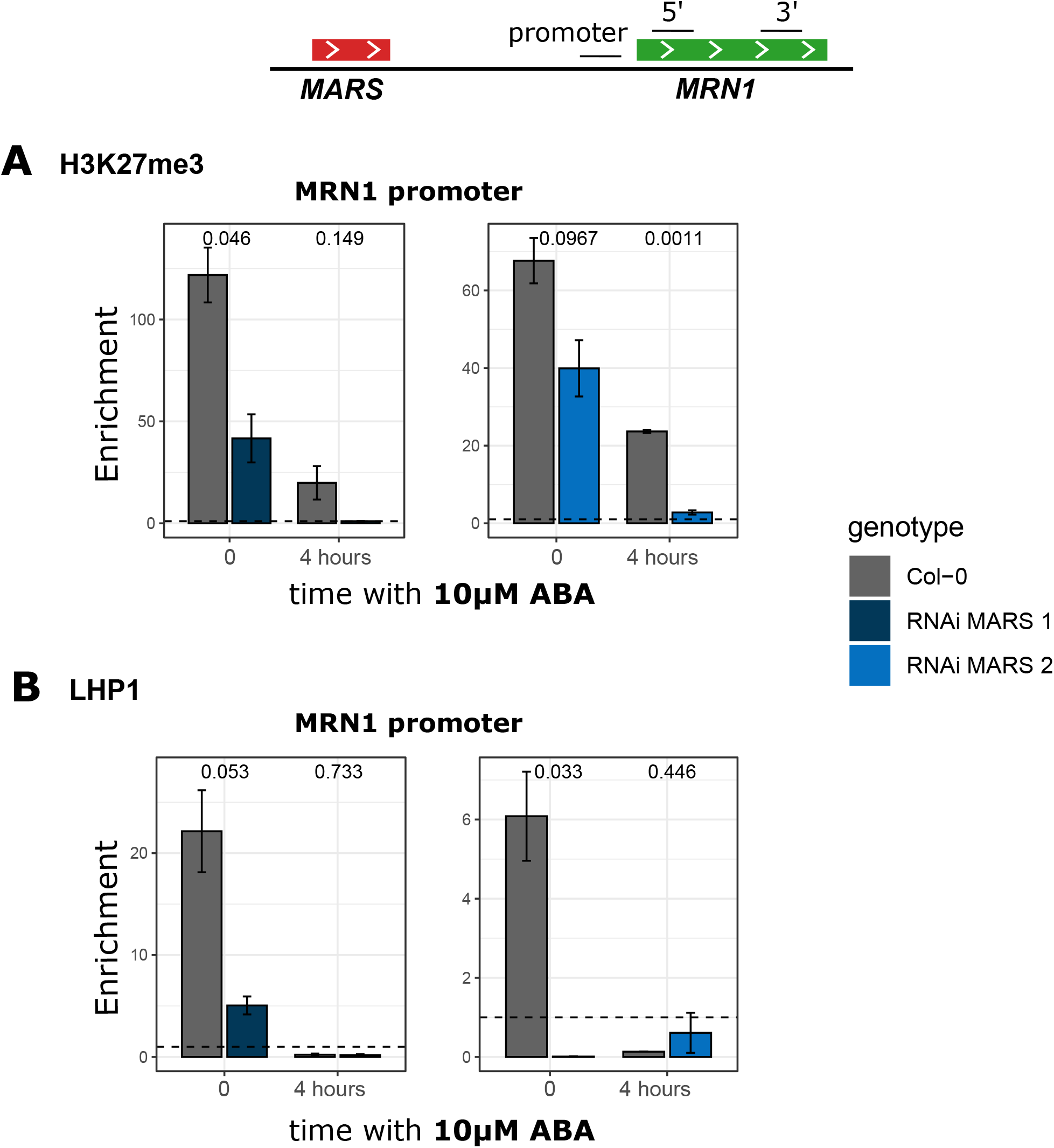
*MARS* modulates the epigenetic landscape of *MRN1* locus. (A) H3K27me3 deposition over the *MRN1* promoter in Col-0 and RNAi-*MARS* seedlings under control conditions and in response to ABA. Higher values of ChIP-qPCR indicate more H3K27me3. (B) LHP1 binding to the *MRN1* promoter in Col-0 and RNAi-*MARS* seedlings in the same conditions as in (A). Higher values of ChIP-qPCR indicate more LHP1 deposition.In (A) and (B), values under the dotted line are considered as not enriched. Results are shown as the mean ± standard error (n = 2) of the H3K27me3/IgG or LHP1/IgG ratio. Numbers are p-value of the difference between the two genotypes determined by Student t-test.

### *MARS* is directly recognized by LHP1 and modulates local chromatin condensation

It has been shown that the deposition of the repressive mark H3K27me3 and the concomitant recognition of the plant PRC1 component LHP1 are correlated with high chromatin condensation (47). Therefore, we determined the chromatin condensation of the whole marneral cluster by Formaldehyde-Assisted Isolation of Regulatory Elements (FAIRE). In contrast to Col-0 showing a highly condensed chromatin, RNAi-*MARS* seedlings exhibit a lower chromatin condensation in control conditions, including the *MARS* locus (**Figure S12**). Notably, the global chromatin status of the cluster was less condensed in RNAi-*MARS* seedlings in response to ABA (**Figure 4A and S12**), in agreement with a decrease of both H3K27me3 deposition and LHP1 binding (**Figure 4A,B, S10 and S11**) and the concomitant transcriptional activation of the clustered genes (**Figure 2B**).

**Figure 4.**
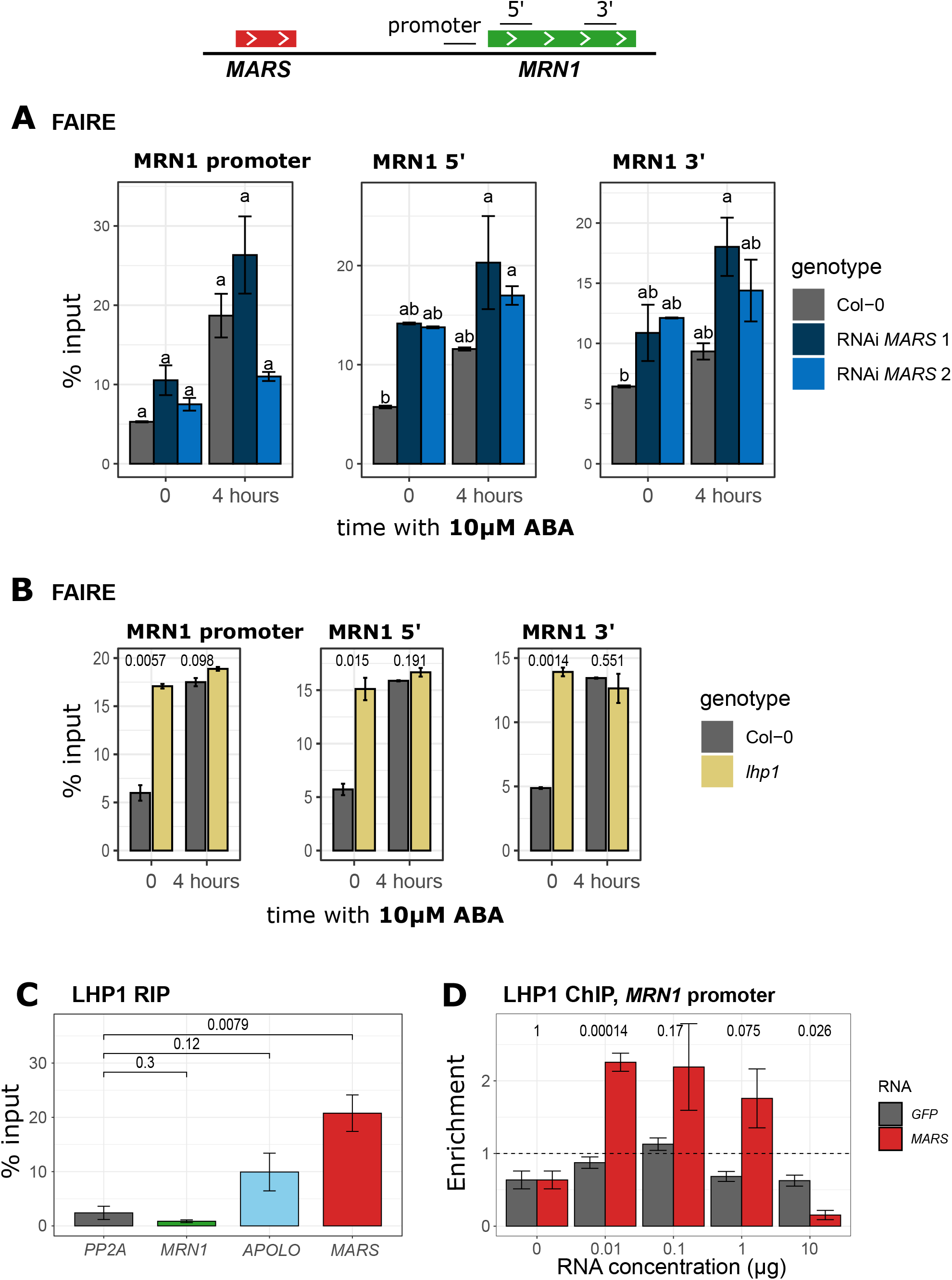
*MARS* influences chromatin condensation of *MRN1* gene through its interaction with LHP1 protein. (A) Chromatin condensation in *MRN1* gene of Col-0 and RNAi-*MARS* seedlings in control conditions and in response to ABA, determined by Formaldehyde Assisted Isolation of Regulatory Element (FAIRE)-qPCR. (B) Evolution of the chromatin condensation in *MRN1* gene of Col-0 and *lhp1* mutant subjected to ABA treatment determined by Formaldehyde Assisted Isolation of Regulatory Element (FAIRE) qPCR. In (A) and (B), results are shown as the mean ± standard error (n = 3) of the percentage of input (signal measured before isolation of decondensed region of chromatin; free of nucleosomes). Lower value indicates more condensed chromatin. Numbers are p-value of the difference between the two genotypes determined by Student t-test. (C) LHP1-*MARS* interaction was assessed by RNA immunoprecipitation (RIP) using *LHP1-GFP* seedlings. Negative controls include a housekeeping gene (*PP2A*) and *MRN1* mRNA. *MRN1* transcript levels in nuclei samples are comparable to *MARS*. The interaction between *APOLO* and LHP1 was taken as a positive control (14). Results are shown as the mean ± standard error (n = 4) of the percentage of input (signal measured before immunoprecipitation). (D) LHP1 binding to the *MRN1* promoter region in chromatin from RNAi-*MARS* 1 seedlings upon increasing amounts of *in-vitro* transcribed *MARS* or *GFP* RNAs. After incubation (see Methods), the samples were crosslinked for LHP1 ChIP-qPCR. Higher values indicate more LHP1-DNA interaction. Results are shown as the mean ± standard error (n = 2) of the LHP1/Igg ratio. In (C) and (D) numbers are p-value of the difference between the different corresponding genes determined by Student t-test.

Consistently, *lhp1* mutant seedlings also showed a global chromatin decondensation in control conditions, comparable to Col-0 in response to ABA. Notably, chromatin decondensation triggered by ABA was completely impaired in *lhp1* (**Figure 4B and S13**), supporting the role of LHP1 in the dynamic epigenetic silencing of the marneral cluster. Concomitantly, the increased chromatin decondensation of *lhp1* mutant seedlings correlates with increased abundance of marneral genes transcripts (**Figure S14A**), as observed in the RNAi-*MARS* seedlings (**Figure 2B**) with decondensed chromatin (**Figure 4A and S12**).

It has been shown that LHP1 can recognize RNAs *in vitro* (12) and the lncRNA *APOLO in vivo* (14). Moreover, it has been proposed that *APOLO* over-accumulation can decoy LHP1 away from target chromatin (48). Therefore, we wondered whether *MARS* lncRNA was able to interact with the chromatin-related protein LHP1 participating in the modulation of the local epigenetic environment. Thus, we first determined that *MARS* was enriched in the nucleus, compared with total RNA, as the previously characterized lncRNAs *APOLO* and *ASCO* that interact respectively with nuclear epigenetic and splicing machineries, and the nuclear structural ncRNA *U6* (**Figure S14B**), involved in the spliceosome. Then, we confirmed by RNA immunoprecipitation (RIP) that LHP1 can interact with *MARS in vivo*, in contrast to *MRN1* or a randomly selected housekeeping gene (*PP2A*) taken as negative controls (**Figure 4C**).

LHP1 binding to the marneral cluster was impaired both in response to exogenous ABA (concomitantly inducing *MARS*, **Figure 3B, S11 and 2B**) and in RNAi-*MARS* seedlings, hinting at a stoichiometry-dependent action of *MARS* on LHP1 recognition of the marneral cluster. Therefore, we used chromatin extracts from RNAi-*MARS* line 1 seedling, that contains very low *MARS* transcript levels (**Figure 2A**) to assess LHP1 recognition of the marneral cluster upon the addition of increasing concentrations of *in vitro*-transcribed *MARS* RNA. Strikingly, we found that low *MARS* RNA concentrations (between 0.01 and 0.1 µg of RNA; **Figure 4D and S14C**) successfully promoted LHP1 binding to the cluster, in contrast to higher concentrations (between 1 and 10 µg of RNA). Moreover, the *in vitro*-transcribed *GFP* RNA was not able to promote LHP1 binding, supporting the relevance of the specific *MARS*-LHP1 stoichiometric interaction for LHP1-target recognition (**Figure 4D and S14C**). Altogether, our results suggest that the physical interaction of the nuclear-enriched lncRNA *MARS* to LHP1 modulates its binding to proximal chromatin in a dual manner likely participating in the modulation of the dynamic chromatin condensation of the marneral cluster.

### *MARS* expression modulates an LHP1-dependent chromatin loop bringing together the *MRN1* locus and an ABA enhancer element

It has been reported that the spatial conformation of cluster-associated domains differs between transcriptionally active and silenced clusters. In Arabidopsis, segregating 3D contacts are distinguished among organs, in agreement with the corresponding transcriptional activity of clustered genes (10). Therefore, we explored whether *MARS* could participate in the dynamic regulation of the local 3D chromatin conformation modulating the transcription of the marneral cluster. According to available HiC datasets (45, 49) there is a significant interaction linking the intergenic region between *CYP71A16* and *MARS* and the *MRN1* locus (indicated as “Chromatin loop” in **Figure 5A**). We used Chromatin Conformation Capture (3C) to monitor the formation of this chromatin loop and found that it increased drastically after 30 min exposure of seedlings to exogenous ABA and that this chromatin loop remained for at least 4 hours after the treatment (**Figure 5B**). These data indicate that the formation of this chromatin loop positively correlates with the transcriptional accumulation of the marneral cluster genes in response to ABA (**Figure 2B**).

**Figure 5.**
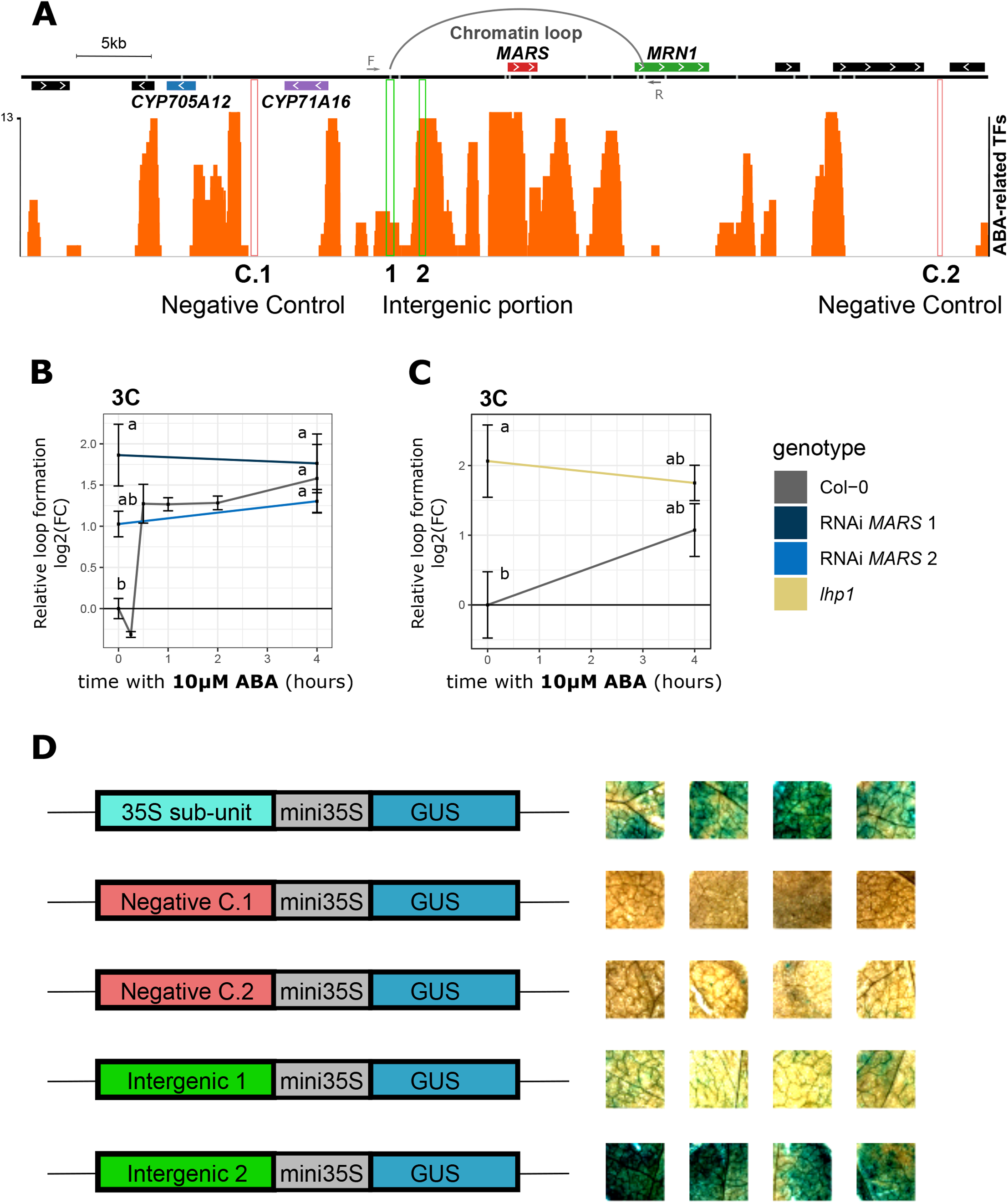
An LHP1-dependent chromatin loop approaches the *MRN1* locus with a putative enhancer element in response to ABA. (A) Schematic illustration of the loop linking the *MRN1* locus with the intergenic region between *CYP71A16* and *MARS*. Forward (F) and Reverse (R) oligonucleotides used for 3C-qPCR (in B–C) are indicated with arrows. The orange track shows the number of different ABA-related transcription factor binding sites (HB6, HB7, GBF2, GBF3, MYB3, MYB44, NF-YC2, NF-YB2, ANAC102, ANAC032, ABF1, ABF3, ABF4, RD26, ZAT6, FBH3, DREBA2A, AT5G04760, HAT22 and HSFA6A) found on the marneral cluster (50). Green and red rectangles indicate the putative enhancer region and the negative controls, respectively, tested for the GUS-based reporter system in (D). (B) Relative chromatin loop formation in response to ABA in Col-0 and RNAi-*MARS* seedlings. Results are shown as the mean ± standard error (n = 2) from 3C-qPCR using primer F and R shown on (A), compared to time 0h. (C) Relative chromatin loop formation in response to ABA treatment in Col-0 and *lhp1* mutant. Data are shown as the mean ± standard error (n = 3) from 3C-qPCR using primer F and R shown on (A), compared to time 0h. (D) Constructs used for the GUS-based reporter system are illustrated on the left. Corresponding transformed tobacco leaf discs are on the right (n = 4). First line represents the positive control in which the 35S sub-unit controls *GUS* expression. The second and third lines show two independent negative controls in which the *GUS* gene is driven by a genomic region that does not contain ABA-related binding sites indicated in (A). In the remaining lines, the transcriptional activity is assessed for the two intergenic regions indicated in (A).

The *MARS* locus is encompassed in the ABA-dependent chromatin loop (**Figure 5A**). In order to determine the role of *MARS* in the modulation of local 3D chromatin conformation, we assessed the formation of the chromatin loop in RNAi-*MARS* lines. Notably, RNAi-*MARS* seedlings exhibit enhanced chromatin loop formation, which remained unchanged in response to exogenous ABA (**Figure 5B**). Interestingly, LHP1 has been implicated in shaping local 3D conformation of target regions (45), suggesting that the LHP1-*MARS* module may dynamically switch the epigenetic status of the marneral cluster from a condensed-linear to a decondensed-3D structured chromatin conformation. Supporting this hypothesis, *lhp1* mutant seedlings exhibited enhanced chromatin loop formation compared to Col-0 (**Figure 5C**). Overall, our results suggest that the formation of a chromatin loop within the marneral cluster is regulated by LHP1 through the interaction with *MARS* lncRNA transcripts.

To better understand the role of the *MARS*-dependent chromatin loop in response to ABA we looked for ABA-related *cis* regulatory sequences throughout the marneral cluster. We analyzed the distribution of binding sites for 13 ABA-related transcription factors (TFs) determined experimentally (50). Interestingly, we found a high enrichment for ABA TF binding sites at the *MARS* locus, as well as in the intergenic region between the *CYP71A16* and *MARS* loci, in particular at regions surrounding the contact point brought into close spatial proximity with the *MRN1* locus by the ABA-dependent 3D chromatin loop (**Figure 5A**). We thus assessed the capacity of these genomic regions to enhance the transcriptional activity of MRN1. To this end, we generated transcriptional reporter line combining the candidate distant enhancer elements to a minimal 35S promoter and ß-glucuronidase (*GUS*) gene (26). We also included as controls two genomic regions nearby the putative enhancers, one between *CYP705A12* and *CYP71A16* and the other at the 3’ end of *AT5G42620* locus (**Figure 5A**). Among the two putative distal enhancers tested, one was able to activate *GUS* expression (Intergenic region 2, **Figure 5D and Figure S15)**, coinciding with the region showing a high enrichment of ABA-related TF binding sites close to the chromatin loop anchor point (**Figure 5A**). Collectively, our results indicate that an ABA-driven chromatin loop brings into close spatial proximity the *MRN1* locus and a transcriptional activation site likely acting as an ABA enhancer element and that this chromatin reorganization process depends on the LHP1-*MARS* module.

### Long noncoding RNAs as emerging regulators of gene clusters

Physically linked genes organized in clusters are generally coregulated (1). Considering that the lncRNA *MARS* is implicated in the regulation of the marneral cluster, we wondered whether the presence of noncoding transcriptional units may constitute a relevant feature of gene cluster organization. Therefore, we looked for the presence of lncRNAs in other gene clusters using two different datasets, one of gene clusters for co-expressed neighboring genes (9) and one for metabolic gene clusters (PlantiSMASH (29)). Among the 390 clusters of co-expressed neighboring genes, 189 (48%) contained at least one lncRNA embedded within the cluster. Most importantly, among the 45 metabolic clusters, 28 (62%) include lncRNAs inside the cluster (**Figure 6A**). Furthermore, among the clusters containing a lncRNA, a correlation analysis based on the maximum strength of co-expression between a lncRNA and any clustered gene revealed that the metabolic clusters exhibit a significantly higher correlation than co-expressed clusters (**Figure 6B**). Altogether, our analyses suggest that lncRNA-mediated local epigenetic remodeling may constitute an emerging feature of non-homologous genes metabolic clusters in plants.

**Figure 6.**
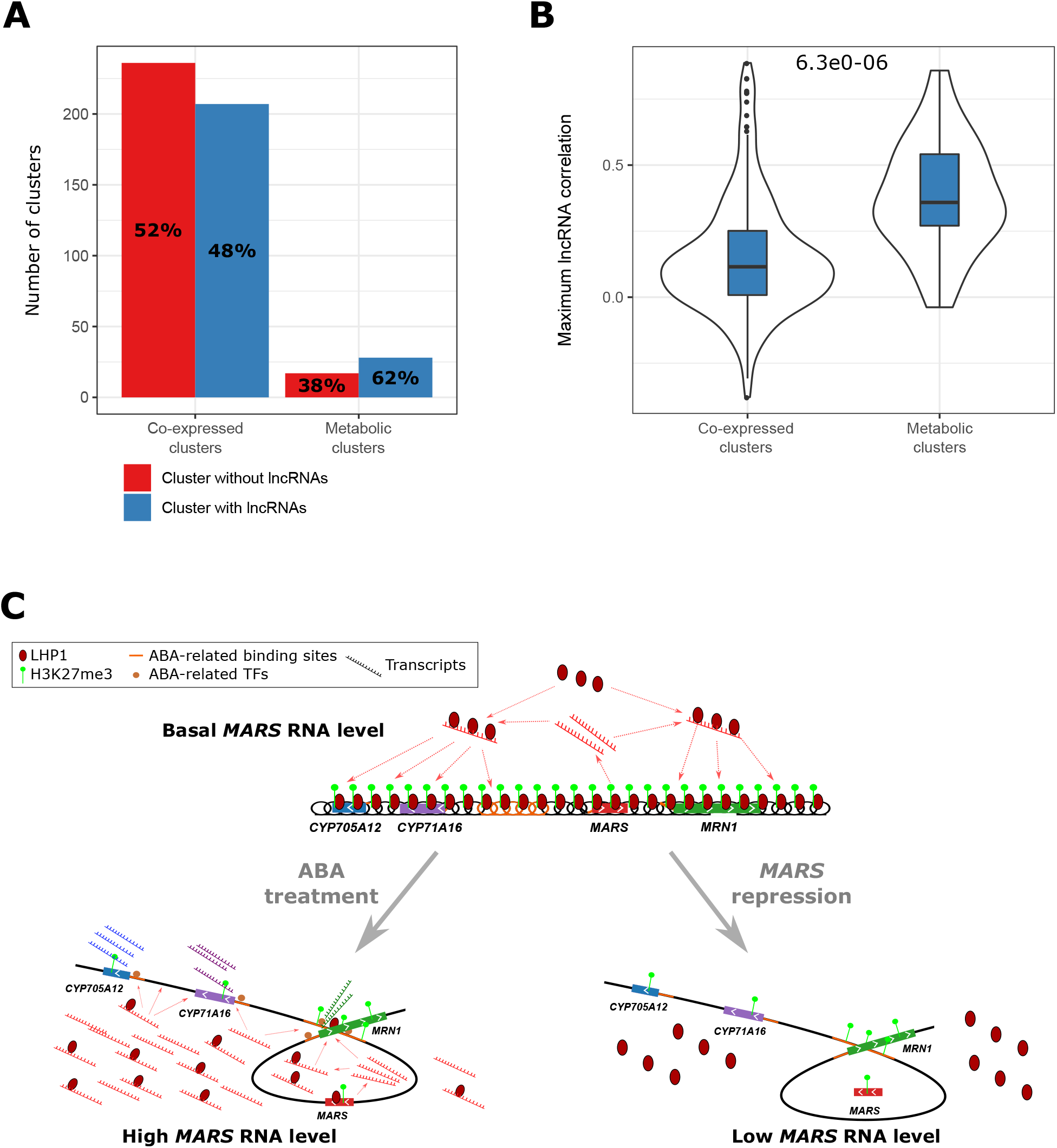
Regulation of metabolic clusters in plants by lncRNA. (A) The proportion of metabolic clusters including lncRNA loci is higher than for other clusters. The co-expressed clusters were predicted in (9) and correspond to co-expressed neighboring genes. The metabolic clusters are co-expressed neighboring genes involved in the biosynthesis of a particular secondary metabolite predicted by plantiSMASH (29). (B) Maximum level of correlation between a lncRNA and any clustered gene determined in each cluster. The number shown above indicates the p-value of the difference between the two type of clusters determined by Student t-test. (C) The lncRNA *MARS* regulates the expression of the marneral cluster genes through epigenetic reprogramming and chromatin conformation. In control conditions (upper panel) the chromatin of the marneral cluster is enriched in H3K27me3 and LHP1, which results in a condensed and linear chromatin conformation. In response to ABA (bottom left panel) *MARS* over-accumulated transcripts titrate LHP1 away from the cluster. The decrease of LHP1 deposition diminishes H3K27me3 distribution, relaxes the chromatin and as a consequence allows the formation of a chromatin loop that approaches the enhancer element and *MRN1* proximal promoter, leading to a transcriptional activation. When *MARS* is repressed, LHP1 recruitment to the cluster is impaired, thus leading to a similar chromatin state: decrease in H3K27me3 mark, chromatin decondensation and increase in chromatin loop conformation. Under this chromatin state, the clustered genes become highly responsive to the ABA treatment.

## DISCUSSION

The cell nucleus is a dynamic arrangement of DNA, RNAs and proteins (51). Genome topology has emerged as an important feature in the complex network of mechanisms regulating gene activity and genome connectivity, leading to regionalized chromosomal spatial distribution and the clustering of diverse genomic regions with similar expression patterns (52).

In the last few years, noncoding transcription has been implicated in shaping 3D nuclear organization (53). Notably, RNase-A micro-injection into the nucleus revealed that long nuclear-retained RNAs maintained euchromatin in a biologically active decondensed state, whereas heterochromatin domains exhibited an RNA-independent structure (54, 55). More recently, HiC analyses were performed in mammalian cells exposed or not to RNase, before and after crosslinking, or upon transcriptional inhibition (56). As a result, it was observed that topologically associated domains (TAD) boundaries remained mostly unaffected by RNase treatment, whereas compartmental interactions suffered a subtle disruption. In contrast, transcriptional inhibition led to weaker TAD boundaries, hinting at different roles of steady-state RNA vs. active transcription in nuclear organization (56).

In plants, several lncRNAs have been implicated in local chromatin conformation dynamics affecting the transcriptional activity of neighboring genes (13, 57). Notably, the lncRNA *COLDWRAP* participates in the formation of an intragenic chromatin loop blocking the transcription of the flowering time regulator *FLOWERING LOCUS C* (*FLC* (15)) in response to cold, in a process involving the recruitment of PRC2 by direct interaction with the component CLF. The lncRNA *APOLO* also controls the transcriptional activity of its neighboring gene *PINOID* (*PID*) by dynamically modulating the formation of an intergenic chromatin loop encompassing the divergent promoter of *PID* and *APOLO* (14), in a process involving the PRC1 component LHP1. More recently, it was proposed that high levels of *APOLO* can decoy LHP1 away from multiple loci in *trans*, modulating the 3D conformation of distal target genes (48). In rice, the expression of the leucine-rich repeat receptor kinase clustered genes *RLKs* is modulated by the locally-encoded lncRNA *LRK ANTISENSE INTERGENIC RNA* (*LAIR*). It was proposed that *LAIR* may directly recruit OsMOF (MALES ABSENT ON THE FIRST) and OsWDR5 (WD REPEAT DOMAIN 5), involved in H4K16 acetylation and chromatin remodeling (58). Here, we showed that the lncRNA *MARS* contributes to the co-regulation of a set of physically linked genes in *cis* in Arabidopsis. We demonstrated that the relative abundance of *in vitro*-transcribed *MARS* fine-tunes LHP1 binding to the cluster region in a stoichiometry-dependent manner, thus explaining how *MARS* levels affect H3K27me3 deposition and chromatin condensation. It has been shown in yeast that histone depletion boosts chromatin flexibility and facilitates chromatin loop formation on the kilobase pair scale (59). In agreement thereof, we uncovered here the dynamic role of the LHP1-*MARS* module affecting nucleosome distribution across the marneral cluster in response to ABA, thus promoting the formation of an intra-cluster chromatin loop.

It has been recently observed that biosynthetic gene clusters are embedded in local three-dimensionally organized hot spots that segregate the region from the surrounding chromosome environment (10). Here, we showed that active noncoding transcriptional units within the cluster may contribute to 3D conformation dynamics switching from silent to active states. Our results indicated that a *MARS*-dependent chromatin loop may bring the *MRN1* locus and a distal ABA-responsive element into close spatial proximity, likely acting as an enhancer. Notably, *MARS*-dependent LHP1 and H3K27me3 removal in Col-0, RNAi-*MARS* and the *lhp1* mutant correlated with chromatin decondensation, loop formation and increased marneral genes transcriptional activity in response to ABA. According to this model, chromatin loop conformation is related to LHP1 binding and is modulated by *MARS* in a dual manner. LHP1 recognition at basal *MARS* levels maintains a possibly linear conformation of the region, precluding the enhancer-*MRN1* locus interaction, whereas the positively activating chromatin loop is formed in the absence of LHP1. *MARS* transcriptional accumulation in response to ABA directly modulates LHP1 binding to the marneral cluster and high levels of *MARS* then titrate LHP1 away from the cluster (**Figure 6C**; in response to ABA). *In vivo*, the ABA-mediated *MARS* transcript accumulation decreases LHP1 binding to the cluster (**Figure 3B and S11**). In agreement, *in vitro*, high level of *MARS* RNA can titrate LHP1 binding even though a minimal amount of *MARS* RNA is required to recruit LHP1 to the marneral locus (**Figure 4D and S14C**). Also, when *MARS* levels are too low compared to basal levels (*in vitro* (**Figure 4D and S14C**) or *in vivo* with the RNAi lines (**Figure 3B and S11**)), recruitment of LHP1 to the cluster is also impaired (**Figure 6C;** *MARS* repression). Thus, the concentration-dependent dual effect of *MARS* on LHP1 binding (**Figure 4D and S14C**) appears as a key factor determining the dynamics of the marneral cluster epigenetic landscape.

In mammals, growing evidence supports the role of lncRNAs in chromatin conformation determination (60) and enhancer activity (e.g. *PVT1* (61) and *CCAT1-L* (62)). Here, we showed that the nuclear-enriched lncRNA *MARS* brings together the *MRN1* proximal promoter and a putative enhancer element enriched in ABA-responsive TF binding sites. Interestingly, it has been shown that human lncRNAs can modulate the binding of TFs to their target chromatin (*DHFR* (63)) and *PANDA* (64), whereas TFs have been implicated in chromatin loop formation in plants (52). Furthermore, it was shown that in addition to the TF NF-YA, the lncRNA *PANDA* interacts with the scaffold-attachment-factor A (SAFA) as well as with PRC1 and PRC2 to modulate cell senescence (65). Therefore, further research will be needed to determine what ABA-responsive TFs are in control of the marneral cluster and to elucidate how they participate in chromatin loop formation along the area, in relation with the PRC1-interacting lncRNA *MARS*.

Plants are a tremendous source of diverse chemicals which are important for their life and survival (9). Marneral biosynthesis has been linked to root and leaf development, flowering time and embryogenesis (2). Here we found that the Arabidopsis marneral cluster is activated by the phytohormone ABA, in a lncRNA-dependent epigenetic reprogramming. *MARS* deregulation affects the cluster response to ABA, impacting seed germination and root sensitivity to osmotic stress. Interestingly, lncRNAs had already been associated with seed germination and environmental stress. For example, the overexpression of the cotton *lncRNA973* resulted in an increased seed germination rate and salt-tolerance in *Arabidopsis* (66). Concomitantly, the decrease in *lncRNA973* transcript abundance in cotton was associated with hypersensitivity to salt stress. In addition, the *Arabidopsis* lncRNA *DRIR* regulates plant response to drought and salt stress by altering the expression of stress-responsive genes (67).

It was proposed that the marneral cluster was founded by the duplication of ancestral genes, independent events of gene rearrangement and the recruitment of additional genes (7). The exploration of the noncoding transcriptome in Arabidopsis recently served to identify ecotype-specific lncRNA-mediated responses to the environment (68). It was suggested that the noncoding genome may participate in multiple mechanisms involved in ecotype adaptation. Collectively, our results indicate that the acquisition of novel noncoding transcriptional units within biosynthetic gene clusters may constitute an additional regulatory layer behind their natural variation in plant evolution.

## AVAILABILITY

Further information and requests for resources and reagents should be directed to and will be fulfilled by the Lead Contact, Martin Crespi.

Plant lines generated in this study are available from the Lead Contact with a completed Materials Transfer Agreement.

## ACKNOWLEDGEMENT

IPS2 benefits from the support of Saclay Plant Sciences-SPS (ANR-17-EUR-0007). We thank Jeremie Bazin and Aurélie Christ from IPS2 for the helpful discussion about results interpretation and design of the experiments. We thank Olivier Martin for critical reading of the manuscript.

## REFERENCES

1. Nützmann, H.W., Huang, A. and Osbourn, A. (2016) Plant metabolic clusters – from genetics to genomics. New Phytol., 211, 771–789.

2. Go, Y.S., Lee, S.B., Kim, H.J., Kim, J., Park, H.Y., Kim, J.K., Shibata, K., Yokota, T., Ohyama, K., Muranaka, T., et al. (2012) Identification of marneral synthase, which is critical for growth and development in Arabidopsis. Plant J., 72, 791–804.

3. Yasumoto, S., Fukushima, E.O., Seki, H. and Muranaka, T. (2016) Novel triterpene oxidizing activity of Arabidopsis thaliana CYP716A subfamily enzymes. FEBS Lett., 590, 533–540.

4. Field, B. and Osbourn, A.E. (2008) Clusters in Different Plants. Science (80-.)., 194, 543–547.

5. Boutanaev, A.M., Moses, T., Zi, J., Nelson, D.R., Mugford, S.T., Peters, R.J. and Osbourn, A. (2015) Investigation of terpene diversification across multiple sequenced plant genomes. Proc. Natl. Acad. Sci. U. S. A., 112, E81–E88.

6. Castillo, D.A., Kolesnikova, M.D. and Matsuda, S.P.T. (2013) An effective strategy for exploring unknown metabolic pathways by genome mining. J. Am. Chem. Soc., 135, 5885–5894.

7. Field, B., Fiston-Lavier, A.-S., Kemen, A., Geisler, K., Quesneville, H. and Osbourn, A.E. (2011) Formation of plant metabolic gene clusters within dynamic chromosomal regions. Proc. Natl. Acad. Sci., 108, 16116–16121.

8. Nützmann, H.W. and Osbourn, A. (2015) Regulation of metabolic gene clusters in Arabidopsis thaliana. New Phytol., 205, 503–510.

9. Yu, N., Nützmann, H.W., Macdonald, J.T., Moore, B., Field, B., Berriri, S., Trick, M., Rosser, S.J., Kumar, S.V., Freemont, P.S., et al. (2016) Delineation of metabolic gene clusters in plant genomes by chromatin signatures. Nucleic Acids Res., 44, 2255–2265.

10. Nützmann, H., Doerr, D., Ramírez-colmenero, A., Sotelo-fonseca, J.E., Fernandez-valverde, S.L. and Osbourn, A. (2020) Active and repressed biosynthetic gene clusters have spatially distinct chromosome states. 10.1073/pnas.1920474117.

11. Rinn, J.L. and Chang, H.Y. (2020) Long Noncoding RNAs: Molecular Modalities to Organismal Functions. Annu. Rev. Biochem., 89, 283–308.

12. Berry, S., Rosa, S., Howard, M., Bühler, M. and Dean, C. (2017) Disruption of an RNA-binding hinge region abolishes LHP1-mediated epigenetic repression. Genes Dev., 31, 2115–2120.

13. Lucero, L., Fonouni-Farde, C., Crespi, M. and Ariel, F. (2020) Long noncoding RNAs shape transcription in plants. Transcription, 00, 1–12.

14. Ariel, F., Jegu, T., Latrasse, D., Romero-Barrios, N., Christ, A., Benhamed, M. and Crespi, M. (2014) Noncoding transcription by alternative rna polymerases dynamically regulates an auxin-driven chromatin loop. Mol. Cell, 55, 383–396.

15. Kim, D.-H. and Sung, S. (2017) Vernalization-triggered intragenic chromatin-loop formation by long noncoding RNAs. Dev. Cell, 176, 100–106.

16. Gagliardi, D., Cambiagno, D.A., Arce, A.L., Tomassi, A.H., Giacomelli, J.I., Ariel, F.D. and Manavella, P.A. (2019) Dynamic regulation of chromatin topology and transcription by inverted repeat-derived small RNAs in sunflower. Proc. Natl. Acad. Sci. U. S. A., 116, 17578–17583.

17. Ariel, F., Brault-Hernandez, M., Laffont, C., Huault, E., Brault, M., Plet, J., Moison, M., Blanchet, S., Ichanté, J.L., Chabaud, M., et al. (2012) Two direct targets of cytokinin signaling regulate symbiotic nodulation in medicago truncatula. Plant Cell, 24, 3838–3852.

18. Lampropoulos, A., Sutikovic, Z., Wenzl, C., Maegele, I., Lohmann, J.U. and Forner, J. (2013) GreenGate - A novel, versatile, and efficient cloning system for plant transgenesis. PLoS One, 8.

19. Clough, S.J. and Bent, A.F. (1998) Floral dip: A simplified method for Agrobacterium-mediated transformation of Arabidopsis thaliana. Plant J., 16, 735–743.

20. Pound, M.P., French, A.P., Atkinson, J.A., Wells, D.M., Bennett, M.J. and Pridmore, T. (2013) RootNav: Navigating Images of Complex Root Architectures. Plant Physiol., 162, 1802–1814.

21. Czechowski, T., Stitt, M., Altmann, T., Udvardi, M.K. and Scheible, W. (2005) Genome-Wide Identification and Testing of Superior Reference Genes for Transcript Normalization in Arabidopsis. Plant Physiol., 139, 5–17.

22. Simon, J.M., Giresi, P.G., Davis, I.J. and Lieb, J.D. (2012) Using formaldehyde-assisted isolation of regulatory elements (FAIRE) to isolate active regulatory DNA. Nat. Protoc., 7, 256–267.

23. Nagymihály, M., Veluchamy, A., Györgypál, Z., Ariel, F., Jégu, T., Benhamed, M., Szücs, A., Kereszt, A., Mergaert, P. and Kondorosi, É. (2017) Ploidy-dependent changes in the epigenome of symbiotic cells correlate with specific patterns of gene expression. Proc. Natl. Acad. Sci. U. S. A., 114, 4543–4548.

24. Nakahigashi, K., Jasencakova, Z., Schubert, I. and Goto, K. (2005) The Arabidopsis HETEROCHROMATIN PROTEIN1 homolog (TERMINAL FLOWER2) silences genes within the euchromatic region but not genes positioned in heterochromatin. Plant Cell Physiol., 46, 1747– 1756.

25. Louwers, M., Splinter, E., van Driel, R., de Laat, W. and Stam, M. (2009) Studying physical chromatin interactions in plants using Chromosome Conformation Capture (3C). Nat. Protoc., 4, 1216– 1229.

26. Yan, W., Chen, D., Schumacher, J., Durantini, D., Engelhorn, J., Chen, M., Carles, C.C. and Kaufmann, K. (2019) Dynamic control of enhancer activity drives stage-specific gene expression during flower morphogenesis. Nat. Commun., 10, 1–16.

27. Moreno-risueno, M.A., Norman, J.M. Van, Moreno, A., Zhang, J., Ahnert, S.E. and Benfey, P.N. (2010) NIH Public Access. 329, 1306–1311.

28. Jefferson, R.A., Kavanagh, T.A. and Bevan, M.W. (1987) GUS fusions:, B-glucuronidase as a sensitive and versatile gene fusion marker in higher plants. EMBO J., 6, 3901–3907.

29. Kautsar, S.A., Suarez Duran, H.G., Blin, K., Osbourn, A. and Medema, M.H. (2017) PlantiSMASH: Automated identification, annotation and expression analysis of plant biosynthetic gene clusters. Nucleic Acids Res., 45, W55–W63.

30. Cheng, C.Y., Krishnakumar, V., Chan, A.P., Thibaud-Nissen, F., Schobel, S. and Town, C.D. (2017) Araport11: a complete reannotation of the Arabidopsis thaliana reference genome. Plant J., 89, 789–804.

31. Dobin, A., Davis, C.A., Schlesinger, F., Drenkow, J., Zaleski, C., Jha, S., Batut, P., Chaisson, M. and Gingeras, T.R. (2013) STAR: Ultrafast universal RNA-seq aligner. Bioinformatics, 29, 15–21.

32. Liao, Y., Smyth, G.K. and Shi, W. (2014) FeatureCounts: An efficient general purpose program for assigning sequence reads to genomic features. Bioinformatics, 30, 923–930.

33. Love, M.I., Huber, W. and Anders, S. (2014) Moderated estimation of fold change and dispersion for RNA-seq data with DESeq2. Genome Biol., 15, 1–21.

34. Wei, T., Simko, V., Levy, M., Xie, Y., Jin, Y. and Zemla, J. (2017) Visualization of a Correlation Matrix. Statistician, 56, 316–324.

35. Wickham, H., Averick, M., Bryan, J., Chang, W., McGowan, L., François, R., Grolemund, G., Hayes, A., Henry, L., Hester, J., et al. (2019) Welcome to the Tidyverse. J. Open Source Softw., 4, 1686.

36. Kong, L., Zhang, Y., Ye, Z.Q., Liu, X.Q., Zhao, S.Q., Wei, L. and Gao, G. (2007) CPC: Assess the protein-coding potential of transcripts using sequence features and support vector machine. Nucleic Acids Res., 35, 345–349.

37. Kang, Y.J., Yang, D.C., Kong, L., Hou, M., Meng, Y.Q., Wei, L. and Gao, G. (2017) CPC2: A fast and accurate coding potential calculator based on sequence intrinsic features. Nucleic Acids Res., 45, W12–W16.

38. Heo, J.B. and Sung, S. (2011) Vernalization-mediated epigenetic silencing by a long intronic noncoding RNA. Science (80-.)., 331, 76–79.

39. Bardou, F., Ariel, F., Simpson, C.G., Romero-Barrios, N., Laporte, P., Balzergue, S., Brown, J.W.S. and Crespi, M. (2014) Long Noncoding RNA Modulates Alternative Splicing Regulators in Arabidopsis. Dev. Cell, 30, 166–176.

40. Jarroux, J., Morillon, A. and Pinskaya, M. (2017) Long Non Coding RNA Biology. Adv. Exp. Med. Biol., 1008, 1–46.

41. Romanowski, A., Schlaen, R.G., Perez-Santangelo, S., Mancini, E. and Yanovsky, M.J. (2020) Global transcriptome analysis reveals circadian control of splicing events in Arabidopsis thaliana. Plant J., 103, 889–902.

42. Covington, M.F., Maloof, J.N., Straume, M., Kay, S.A. and Harmer, S.L. (2008) Global transcriptome analysis reveals circadian regulation of key pathways in plant growth and development. Genome Biol., 9.

43. Hsu, P.Y. and Harmer, S.L. (2012) Circadian Phase Has Profound Effects on Differential Expression Analysis. PLoS One, 7, 18–21.

44. Vishwakarma, K., Upadhyay, N., Kumar, N., Yadav, G., Singh, J., Mishra, R.K., Kumar, V., Verma, R., Upadhyay, R.G., Pandey, M., et al. (2017) Abscisic acid signaling and abiotic stress tolerance in plants: A review on current knowledge and future prospects. Front. Plant Sci., 8, 1–12.

45. Veluchamy, A., Jégu, T., Ariel, F., Latrasse, D., Mariappan, K.G., Kim, S.K., Crespi, M., Hirt, H., Bergounioux, C., Raynaud, C., et al. (2016) LHP1 Regulates H3K27me3 Spreading and Shapes the Three-Dimensional Conformation of the Arabidopsis Genome. PLoS One, 11, 1–25.

46. Sijacic, P., Bajic, M., McKinney, E.C., Meagher, R.B. and Deal, R.B. (2018) Changes in chromatin accessibility between Arabidopsis stem cells and mesophyll cells illuminate cell type-specific transcription factor networks. Plant J., 94, 215–231.

47. Yang, X., Tong, A., Yan, B. and Wang, X. (2017) Governing the silencing state of chromatin: The roles of polycomb repressive complex 1 in arabidopsis. Plant Cell Physiol., 58, 198–206.

48. Ariel, F., Lucero, L., Christ, A., Mammarella, M.F., Jegu, T., Veluchamy, A., Mariappan, K., Latrasse, D., Blein, T., Liu, C., et al. (2020) R-Loop Mediated trans Action of the APOLO Long Noncoding RNA. Mol. Cell, 77, 1–11.

49. Liu, C., Wang, C., Wang, G., Becker, C., Zaidem, M. and Weigel, D. (2016) Genome-wide analysis of chromatin packing in Arabidopsis thaliana at single-gene resolution. Genome Res., 26, 1057– 1068.

50. Song, L., Huang, S.S.C., Wise, A., Castanoz, R., Nery, J.R., Chen, H., Watanabe, M., Thomas, J., Bar-Joseph, Z. and Ecker, J.R. (2016) A transcription factor hierarchy defines an environmental stress response network. Science (80-.)., 354, 598.

51. Cavalli, G. and Misteli, T. (2013) Functional implications of genome topology. Nat. Struct. Mol. Biol., 20, 290–299.

52. Rodriguez-Granados, N.Y., Ramirez-Prado, J.S., Veluchamy, A., Latrasse, D., Raynaud, C., Crespi, M., Ariel, F. and Benhamed, M. (2016) Put your 3D glasses on: Plant chromatin is on show. J. Exp. Bot., 67, 3205–3221.

53. Quinodoz, S. and Guttman, M. (2014) Long non-coding RNAs: An emerging link between gene regulation and nuclear organization. Trends Cell Biol., 24, 651–663.

54. Caudron-Herger, M., Müller-Ott, K., Mallm, J.P., Marth, C., Schmidt, U., Fejes-Tóth, K. and Rippe, K. (2011) Coding RNAs with a non-coding function: Maintenance of open chromatin structure. Nucleus, 2.

55. Caudron-Herger, M. and Rippe, K. (2012) Nuclear architecture by RNA. Curr. Opin. Genet. Dev., 22, 179–187.

56. Barutcu, A.R., Blencowe, B.J. and Rinn, J.L. (2019) Differential contribution of steady-state RNA and active transcription in chromatin organization. EMBO Rep., 20, 1–13.

57. Gagliardi, D. and Manavella, P.A. (2020) Short-range regulatory chromatin loops in plants. New Phytol., 10.1111/nph.16632.

58. Wang, Y., Luo, X., Sun, F., Hu, J., Zha, X., Su, W. and Yang, J. (2018) Overexpressing lncRNA LAIR increases grain yield and regulates neighbouring gene cluster expression in rice. Nat. Commun., 9, 1–9.

59. Diesinger, P.M., Kunkel, S., Langowski, J. and Heermann, D.W. (2010) Histone depletion facilitates chromatin loops on the kilobasepair scale. Biophys. J., 99, 2995–3001.

60. Gil, N. and Ulitsky, I. (2020) Regulation of gene expression by cis-acting long non-coding RNAs. Nat. Rev. Genet., 21, 102–117.

61. Cho, S.W., Xu, J., Sun, R., Mumbach, M.R., Carter, A.C., Chen, Y.G., Yost, K.E., Kim, J., He, J., Nevins, S.A., et al. (2018) Promoter of lncRNA Gene PVT1 Is a Tumor-Suppressor DNA Boundary Element. Cell, 173, 1398-1412.e22.

62. Xiang, J.F., Yin, Q.F., Chen, T., Zhang, Y., Zhang, X.O., Wu, Z., Zhang, S., Wang, H. Bin, Ge, J., Lu, X., et al. (2014) Human colorectal cancer-specific CCAT1-L lncRNA regulates long-range chromatin interactions at the MYC locus. Cell Res., 24, 513–531.

63. Martianov, I., Ramadass, A., Serra Barros, A., Chow, N. and Akoulitchev, A. (2007) Repression of the human dihydrofolate reductase gene by a non-coding interfering transcript. Nature, 445, 666– 670.

64. Hung, T., Wang, Y., Lin, M.F., Koegel, A.K., Kotake, Y., Grant, G.D., Horlings, H.M., Shah, N., Umbricht, C., Wang, P., et al. (2011) Extensive and coordinated transcription of noncoding RNAs within cell-cycle promoters. Nat. Genet., 43, 621–629.

65. Puvvula, P.K., Desetty, R.D., Pineau, P., Marchio, A., Moon, A., Dejean, A. and Bischof, O. (2014) Long noncoding RNA PANDA and scaffold-attachment-factor SAFA control senescence entry and exit. Nat. Commun., 5.

66. Zhang, X., Dong, J., Deng, F., Wang, W., Cheng, Y., Song, L., Hu, M., Shen, J., Xu, Q. and Shen, F. (2019) The long non-coding RNA lncRNA973 is involved in cotton response to salt stress. BMC Plant Biol., 19, 459.

67. Qin, T., Zhao, H., Cui, P., Albesher, N. and Xionga, L. (2017) A nucleus-localized long non-coding rna enhances drought and salt stress tolerance. Plant Physiol., 175, 1321–1336.

68. Blein, T., Balzergue, C., Roulé, T., Gabriel, M., Scalisi, L., François, T., Sorin, C., Christ, A., Godon, C., Delannoy, E., et al. (2020) Landscape of the non-coding transcriptome response of two Arabidopsis ecotypes to phosphate starvation. Plant Physiol., 183, pp.00446.2020.

