## Supplemental for "The lncRNA *MARS* modulates the epigenetic reprogramming of the marneral cluster in response to ABA"

### FigureS1

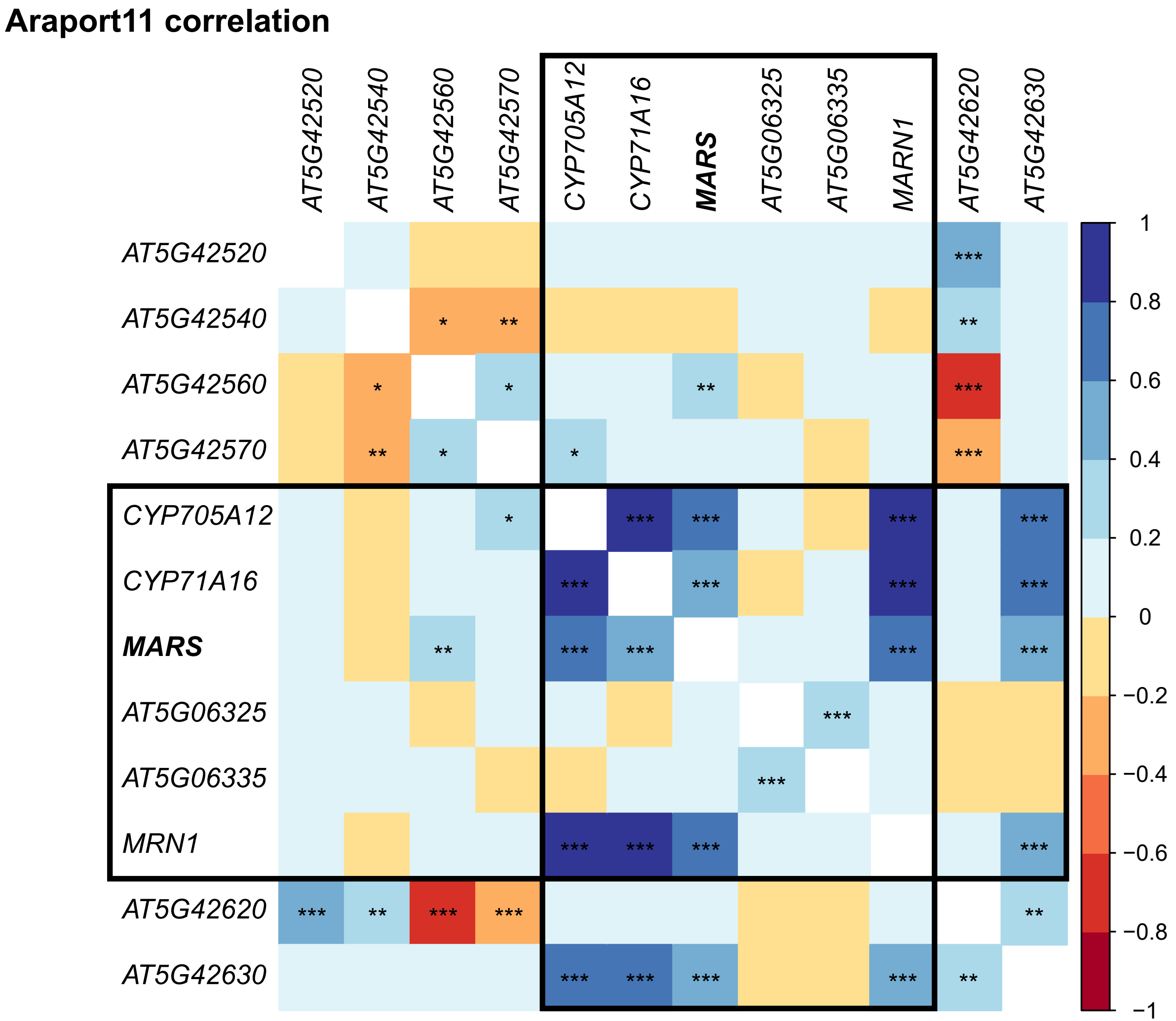
Araport11 correlation
AT5G42520
AT5G42540
AT5G42560
AT5G42570
CYP705A12
CYP71A16
MARS
AT5G06325
AT5G06335
MARN1
AT5G42620
AT5G42630
MRN1
1
0.8
0.6
0.4
0.2
0
−0.2
−0.4
−0.6
−0.8
−1

### FigureS2

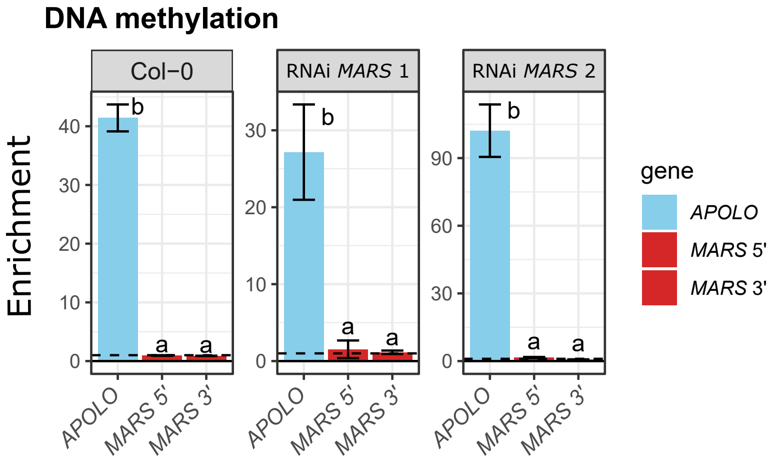
DNA methylation
Enrichment
Col−0
RNAi MARS 1
RNAi MARS 2
0
10
20
30
40
90
60
b
a
APOLO
MARS 5'
MARS 3'
gene

### FigureS3

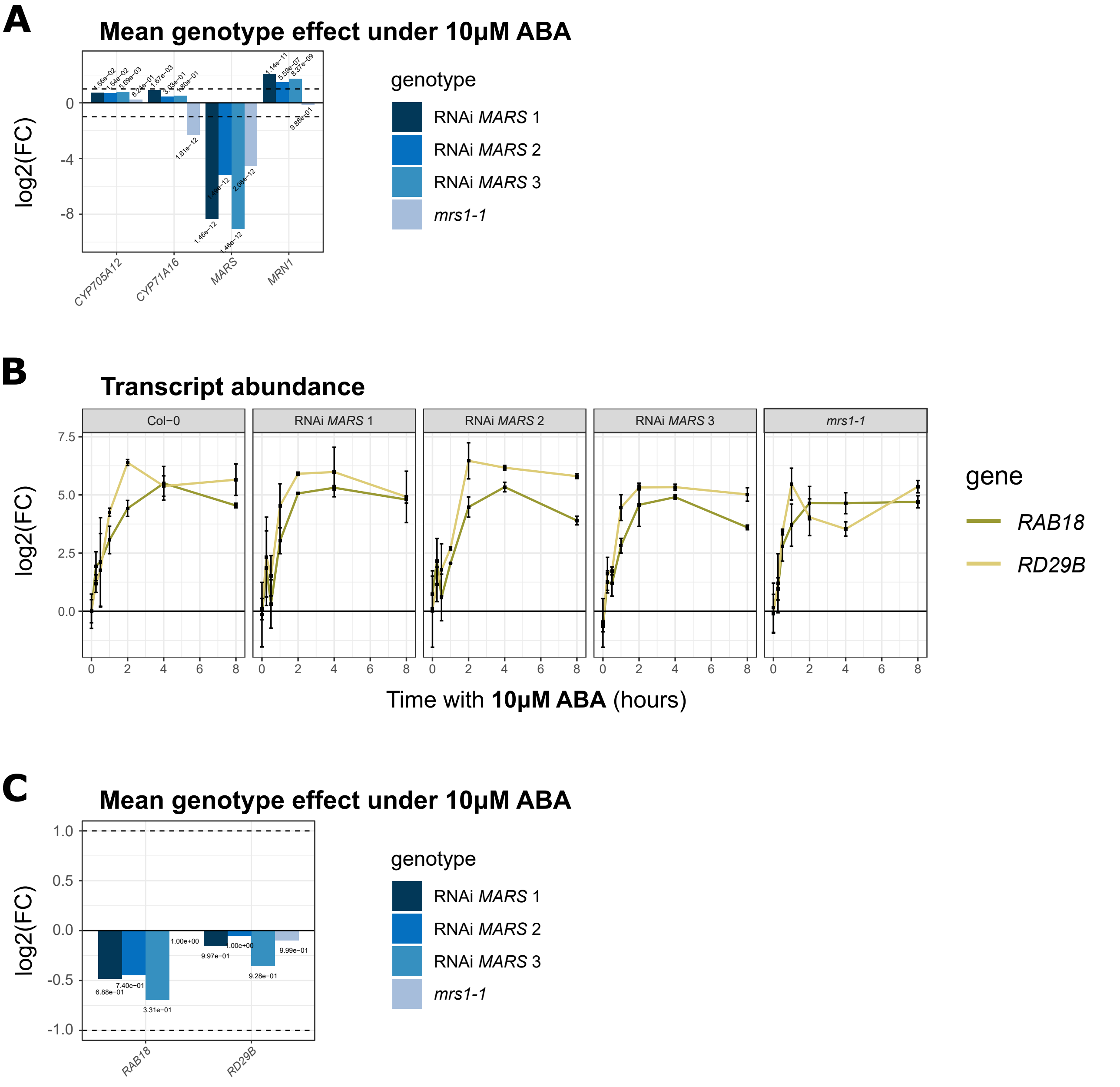
A
Mean genotype effect under 10µM ABA
log2(FC)
0
-4
-8
CYP705A12
CYP71A16
MARS
MRN1
genotype
RNAi MARS 1
RNAi MARS 2
RNAi MARS 3
mrs1-1
B
Transcript abundance
Col−0
RNAi MARS 1
RNAi MARS 2
RNAi MARS 3
mrs1-1
log2(FC)
7.5
5.0
2.5
0.0
0
2
4
6
8
Time with 10µM ABA (hours)
gene
RAB18
RD29B
C
Mean genotype effect under 10µM ABA
log2(FC)
1.0
0.5
0.0
-0.5
-1.0
6.88e−01
7.40e−01
3.31e−01
1.00e+00
9.97e−01
1.00e+00
9.28e−01
9.99e−01
RAB18
RD29B
genotype
RNAi MARS 1
RNAi MARS 2
RNAi MARS 3
mrs1-1

### FigureS4

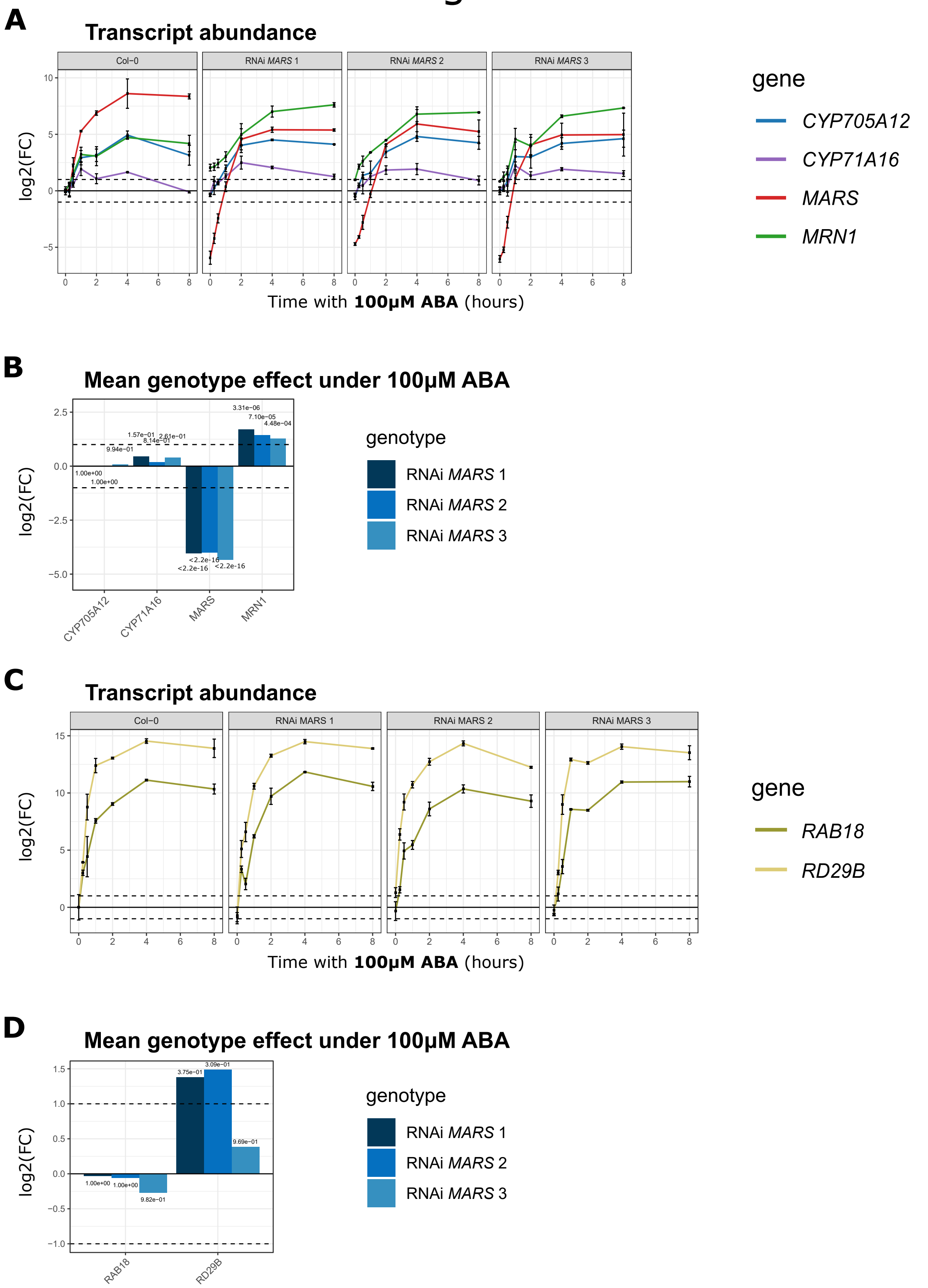
A
Transcript abundance
Col−0
RNAi MARS 1
RNAi MARS 2
RNAi MARS 3
log2(FC)
Time with 100µM ABA (hours)
gene
CYP705A12
CYP71A16
MARS
MRN1
B
Mean genotype effect under 100µM ABA
log2(FC)
3.31e−06
7.10e−05
4.48e−04
1.57e−01
2.61e−01
8.14e−01
9.94e−01
1.00e+00
1.00e+00
<2.2e−16
<2.2e−16
<2.2e−16
CYP705A12
CYP71A16
MARS
MRN1
genotype
RNAi MARS 1
RNAi MARS 2
RNAi MARS 3
C
Transcript abundance
Col−0
RNAi MARS 1
RNAi MARS 2
RNAi MARS 3
log2(FC)
Time with 100µM ABA (hours)
gene
RAB18
RD29B
D
Mean genotype effect under 100µM ABA
log2(FC)
3.75e−01
3.09e−01
9.69e−01
1.00e+00
1.00e+00
9.82e−01
RAB18
RD29B
genotype
RNAi MARS 1
RNAi MARS 2
RNAi MARS 3

### FigureS5

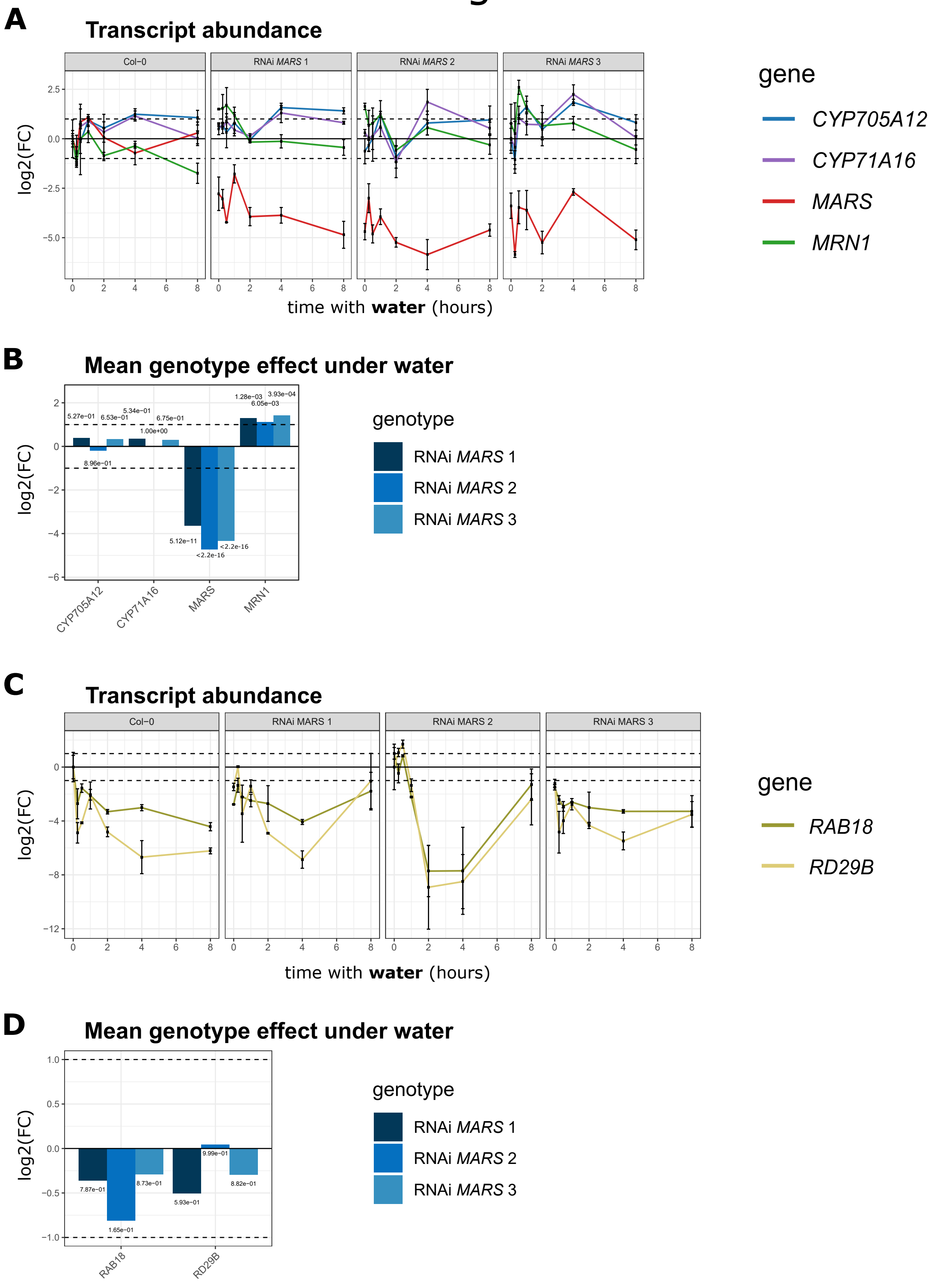

A
Transcript abundance
Col−0
RNAi MARS 1
RNAi MARS 2
RNAi MARS 3
log2(FC)
2.5
0.0
−2.5
−5.0
0
2
4
6
8
time with water (hours)
gene
CYP705A12
CYP71A16
MARS
MRN1
B
Mean genotype effect under water
log2(FC)
2
0
−2
−4
−6
5.27e−01
6.53e−01
5.34e−01
6.75e−01
1.00e+00
8.96e−01
1.28e−03
3.93e−04
6.05e−03
5.12e−11
<2.2e-16
<2.2e-16
CYP705A12
CYP71A16
MARS
MRN1
genotype
RNAi MARS 1
RNAi MARS 2
RNAi MARS 3
C
Transcript abundance
Col−0
RNAi MARS 1
RNAi MARS 2
RNAi MARS 3
log2(FC)
0
−4
−8
−12
0
2
4
6
8
time with water (hours)
gene
RAB18
RD29B
D
Mean genotype effect under water
log2(FC)
1.0
0.5
0.0
−0.5
−1.0
7.87e−01
1.65e−01
8.73e−01
5.93e−01
9.99e−01
8.82e−01
RAB18
RD29B
genotype
RNAi MARS 1
RNAi MARS 2
RNAi MARS 3

### FigureS6

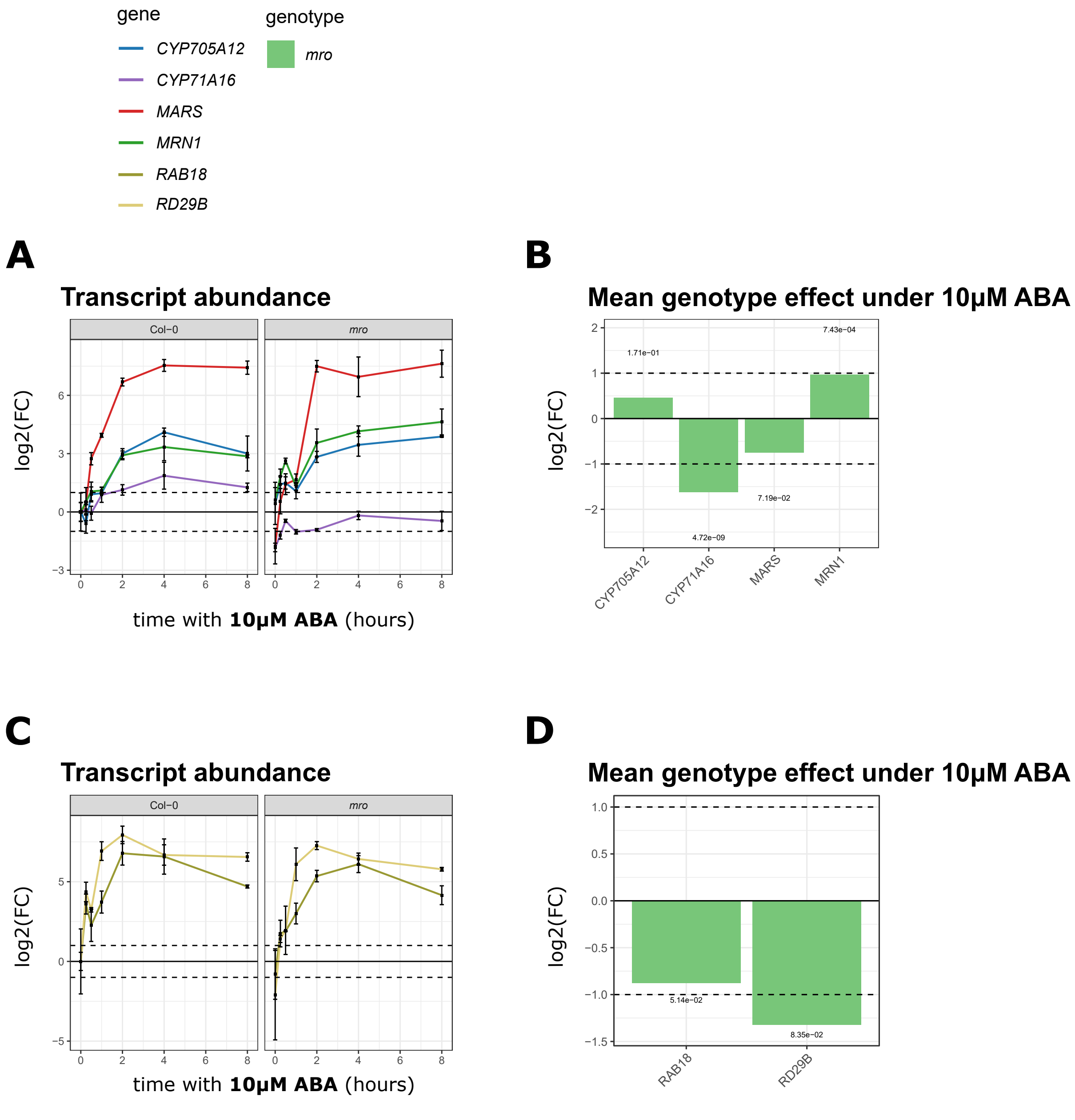
gene
CYP705A12
CYP71A16
MARS
MRN1
RAB18
RD29B
genotype
mro
A
Transcript abundance
Col−0
mro
log2(FC)
time with 10μM ABA (hours)
B
Mean genotype effect under 10μM ABA
log2(FC)
1.71e−01
4.72e−09
7.19e−02
7.43e−04
CYP705A12
CYP71A16
MARS
MRN1
C
Transcript abundance
Col−0
mro
log2(FC)
time with 10μM ABA (hours)
D
Mean genotype effect under 10μM ABA
log2(FC)
5.14e−02
8.35e−02
RAB18
RD29B

### FigureS7

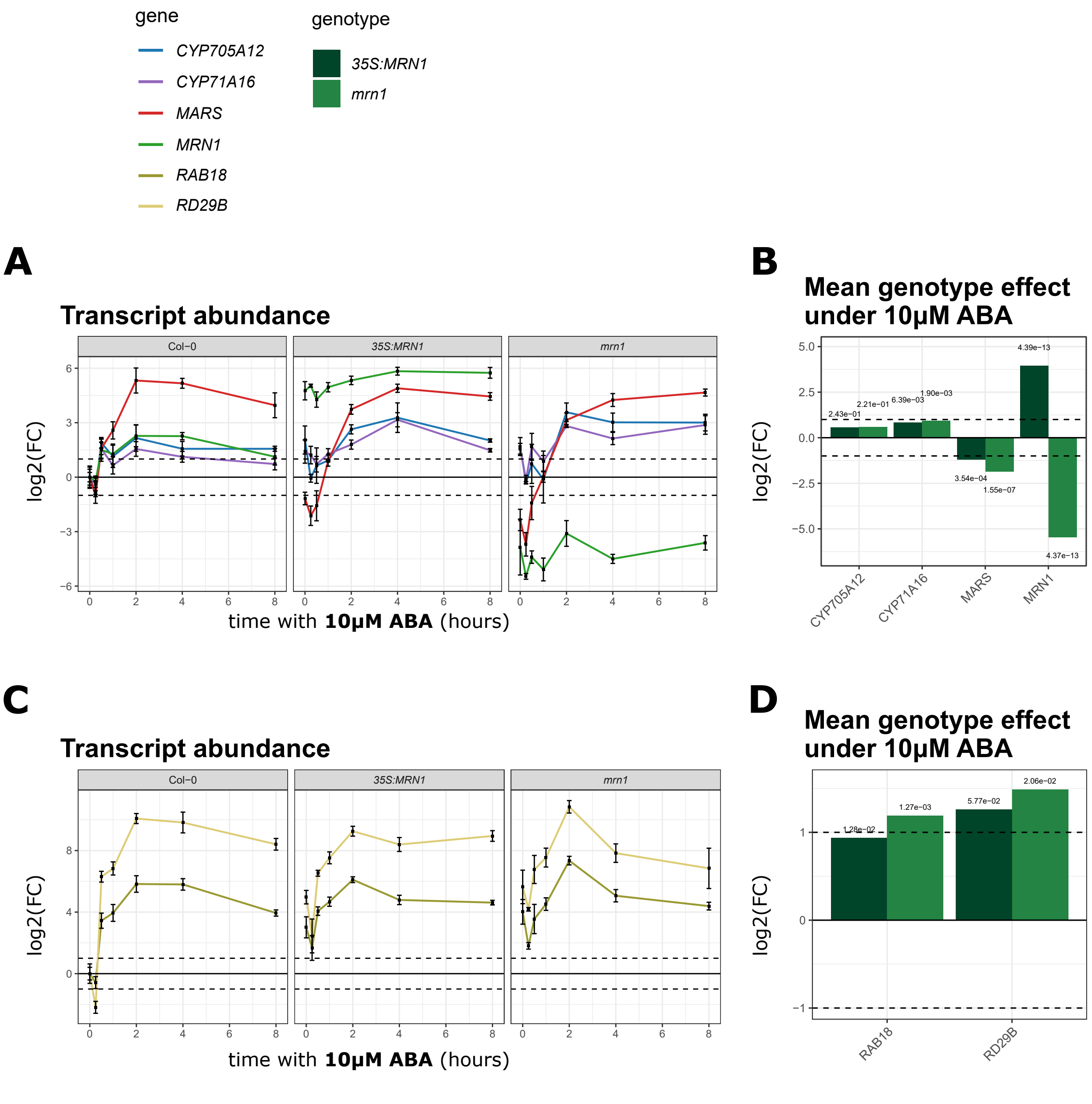
gene
CYP705A12
CYP71A16
MARS
MRN1
RAB18
RD29B
genotype
35S:MRN1
mrn1
A
Transcript abundance
Col−0
35S:MRN1
mrn1
log2(FC)
time with 10µM ABA (hours)
B
Mean genotype effect under 10µM ABA
2.43e−01
2.21e−01
6.39e−03
1.90e−03
4.39e−13
3.54e−04
1.55e−07
4.37e−13
CYP705A12
CYP71A16
MARS
MRN1
C
Transcript abundance
D
Mean genotype effect under 10µM ABA
1.28e−02
1.27e−03
5.77e−02
2.06e−02
RAB18
RD29B

### FigureS8

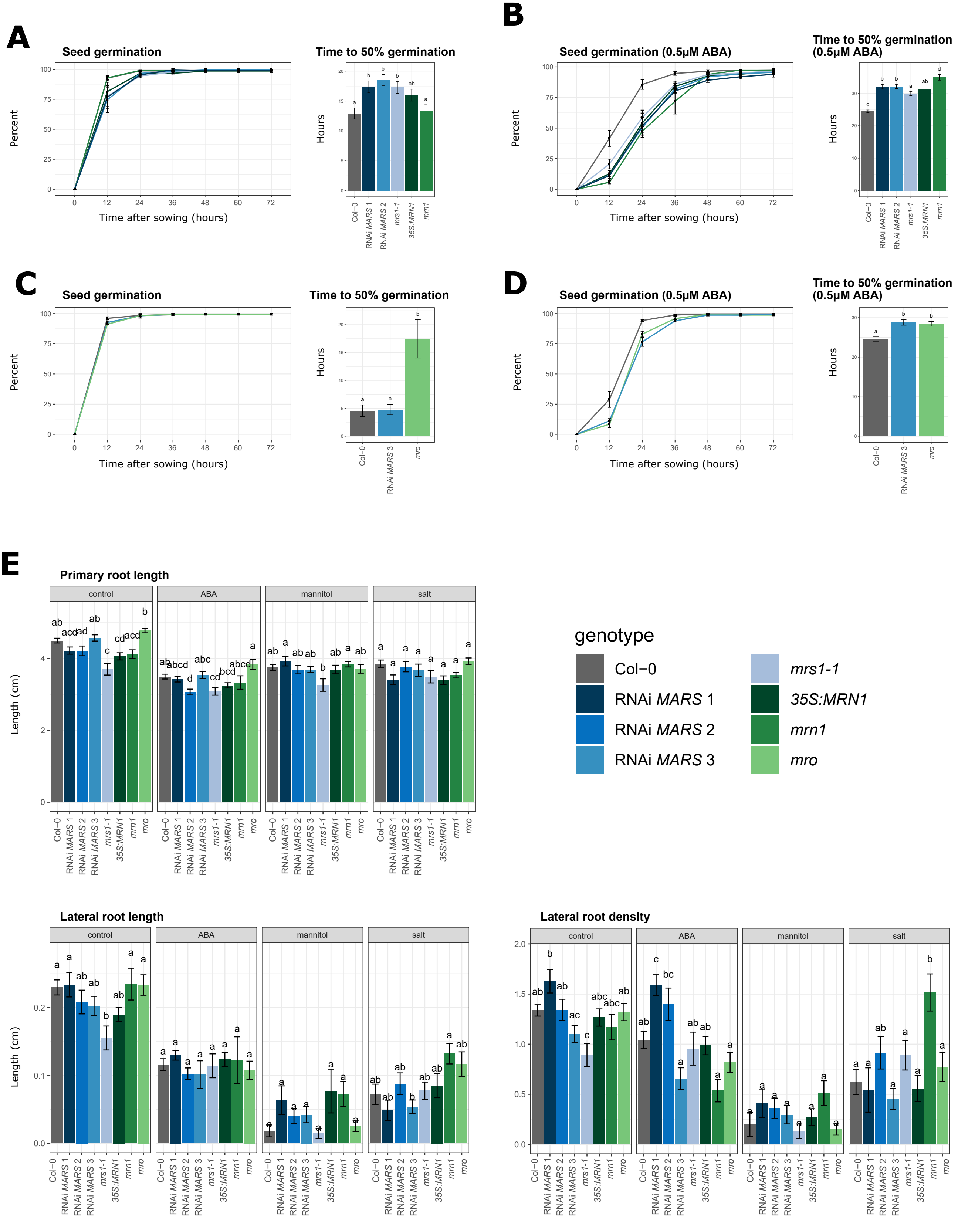
A
Seed germination
Percent
Time after sowing (hours)
Time to 50% germination
Hours
B
Seed germination (0.5μM ABA)
Time to 50% germination (0.5μM ABA)
C
D
E
Primary root length
control
ABA
mannitol
salt
Length (cm)
Lateral root length
Lateral root density
genotype
Col−0
RNAi MARS 1
RNAi MARS 2
RNAi MARS 3
mrs1-1
35S:MRN1
mrn1
mro

### FigureS9

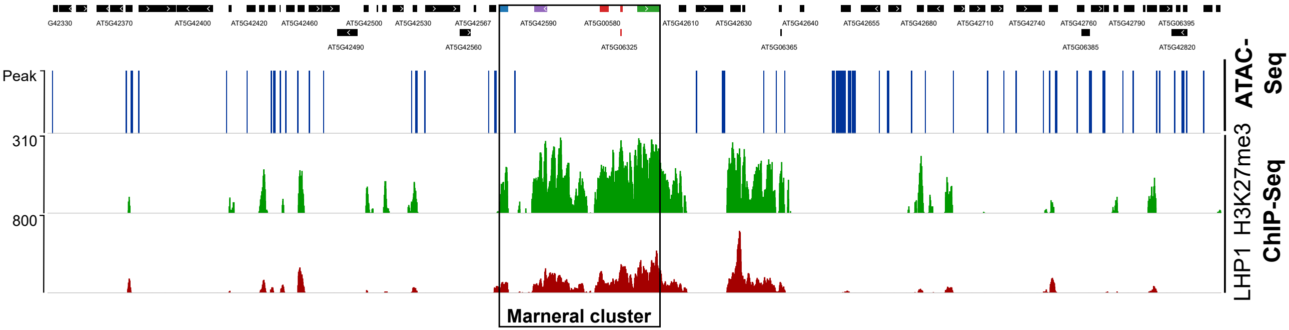

G42330
AT5G42370
AT5G42400
AT5G42420
AT5G42460
AT5G42500
AT5G42530
AT5G42567
AT5G42590
AT5G00580
AT5G42610
AT5G42630
AT5G42640
AT5G42655
AT5G42680
AT5G42710
AT5G42740
AT5G42760
AT5G42790
AT5G06395
AT5G42490
AT5G42560
AT5G06325
AT5G06365
AT5G06385
AT5G42820
Peak
310
800
ATAC-Seq
H3K27me3
LHP1
ChIP-Seq
Marneral cluster

### FigureS10

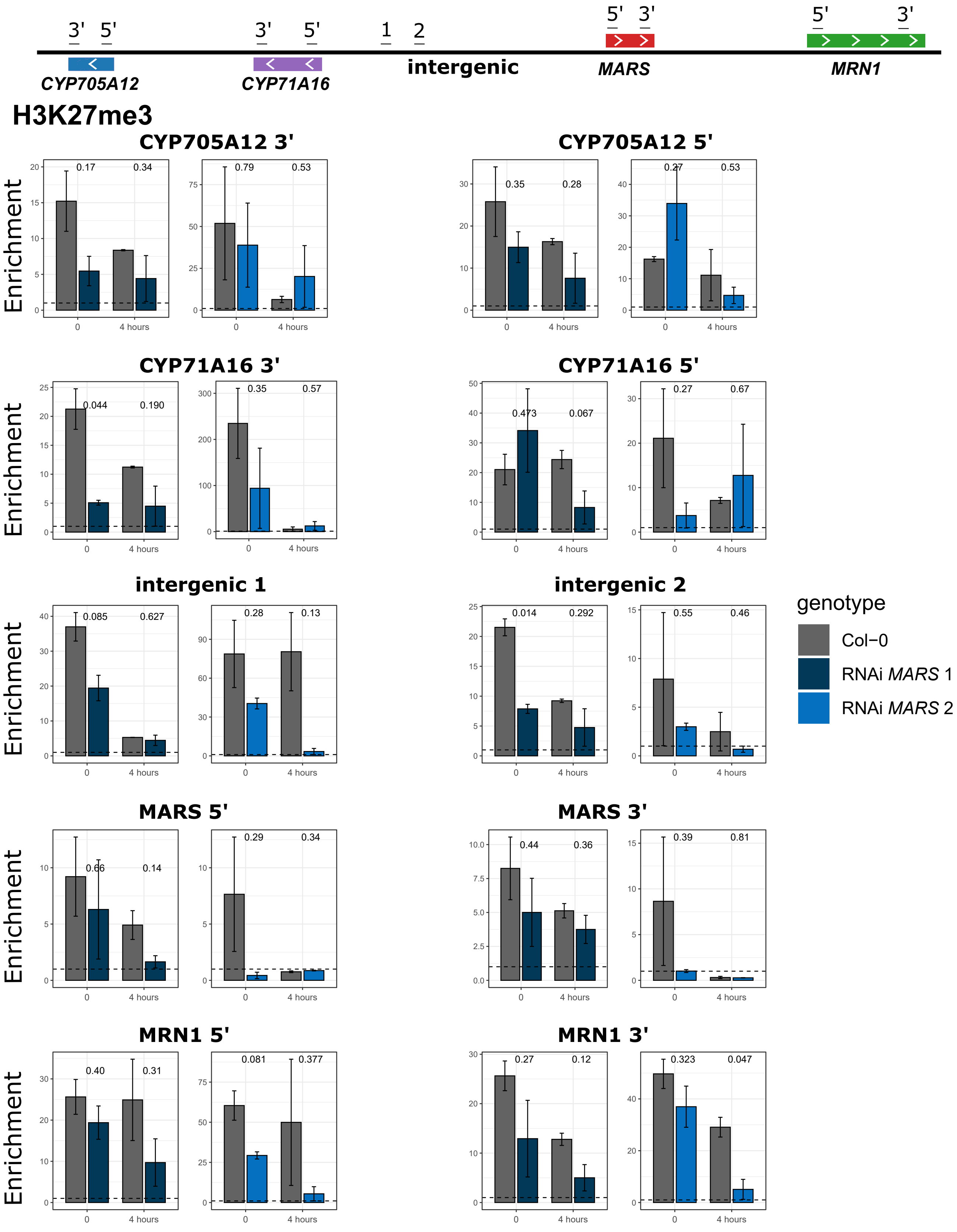

3' 5'
3' 5'
1 2
5' 3'
5' 3'
CYP705A12
CYP71A16
intergenic
MARS
MRN1
H3K27me3
CYP705A12 3'
CYP705A12 5'
CYP71A16 3'
CYP71A16 5'
intergenic 1
intergenic 2
MARS 5'
MARS 3'
MRN1 5'
MRN1 3'
Enrichment
0
4 hours
0.17
0.34
0.79
0.53
0.35
0.28
0.27
0.53
0.044
0.190
0.35
0.57
0.473
0.067
0.27
0.67
0.085
0.627
0.28
0.13
0.014
0.292
0.55
0.46
0.66
0.14
0.29
0.34
0.44
0.36
0.39
0.81
0.40
0.31
0.081
0.377
0.27
0.12
0.323
0.047
genotype
Col−0
RNAi MARS 1
RNAi MARS 2

### FigureS11

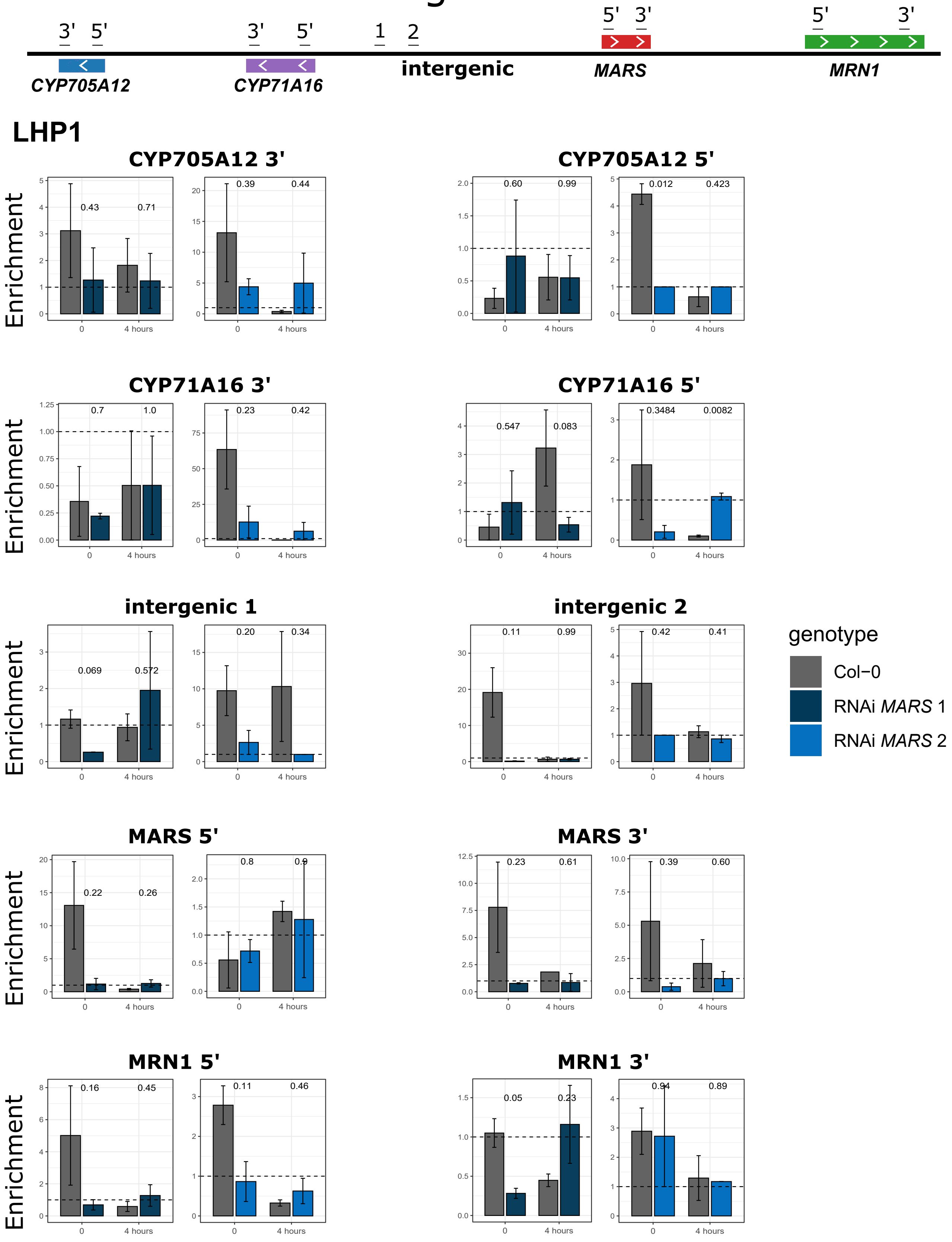
3' 5'
CYP705A12
3' 5'
CYP71A16
1 2
intergenic
5' 3'
MARS
5' 3'
MRN1
LHP1
CYP705A12 3'
0.43 0.71
0.39 0.44
CYP705A12 5'
0.60 0.99
0.012 0.423
CYP71A16 3'
0.7 1.0
0.23 0.42
CYP71A16 5'
0.547 0.083
0.3484 0.0082
intergenic 1
0.069 0.572
0.20 0.34
intergenic 2
0.11 0.99
0.42 0.41
MARS 5'
0.22 0.26
0.8 0.9
MARS 3'
0.23 0.61
0.39 0.60
MRN1 5'
0.16 0.45
0.11 0.46
MRN1 3'
0.05 0.23
0.94 0.89
Enrichment
0
4 hours
genotype
Col−0
RNAi MARS 1
RNAi MARS 2

### FigureS12

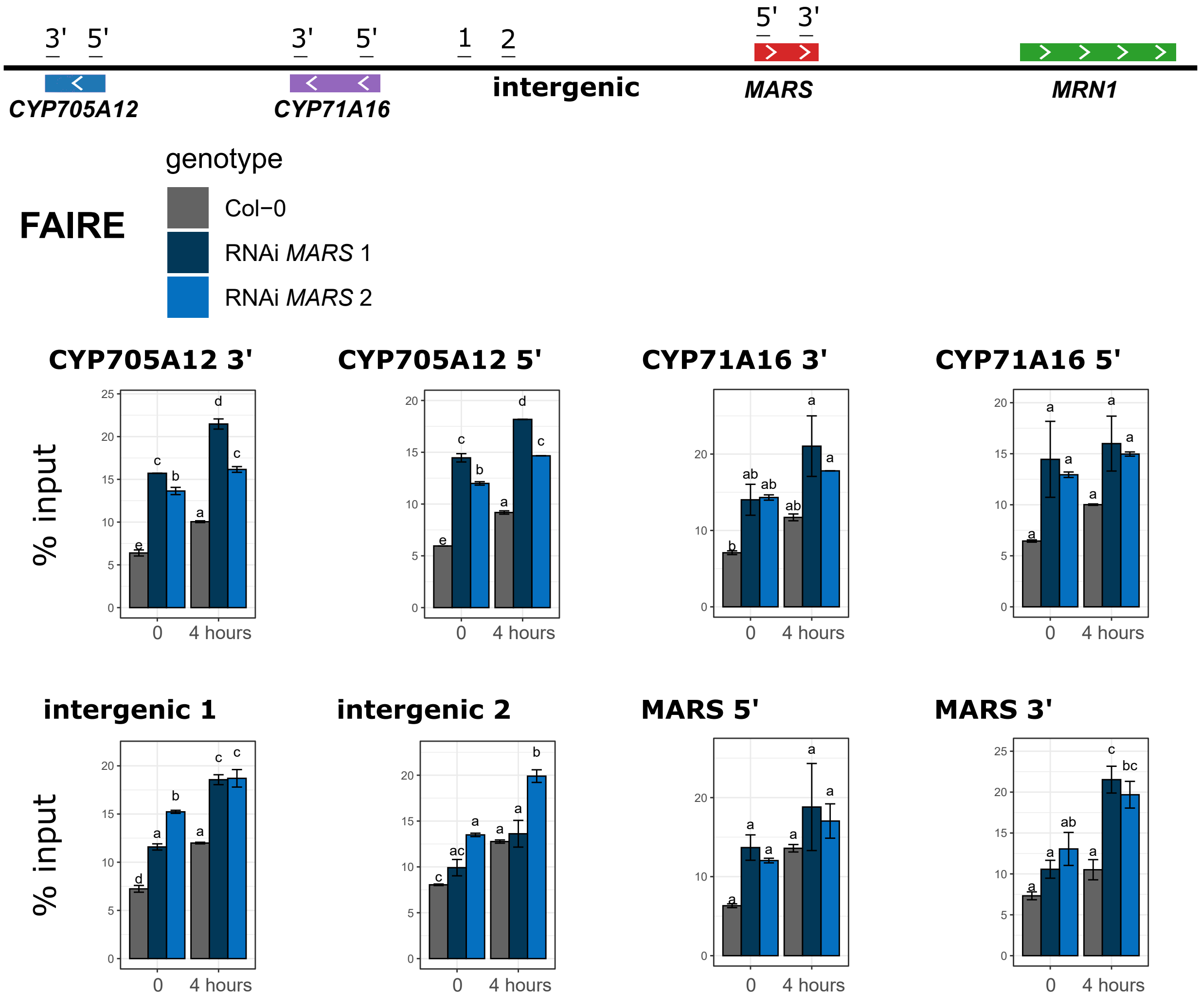
3'
5'
CYP705A12
3'
5'
CYP71A16
1
2
intergenic
5'
3'
MARS
MRN1
FAIRE
genotype
Col−0
RNAi MARS 1
RNAi MARS 2
CYP705A12 3'
CYP705A12 5'
CYP71A16 3'
CYP71A16 5'
intergenic 1
intergenic 2
MARS 5'
MARS 3'
% input
% input
0
4 hours

### FigureS13

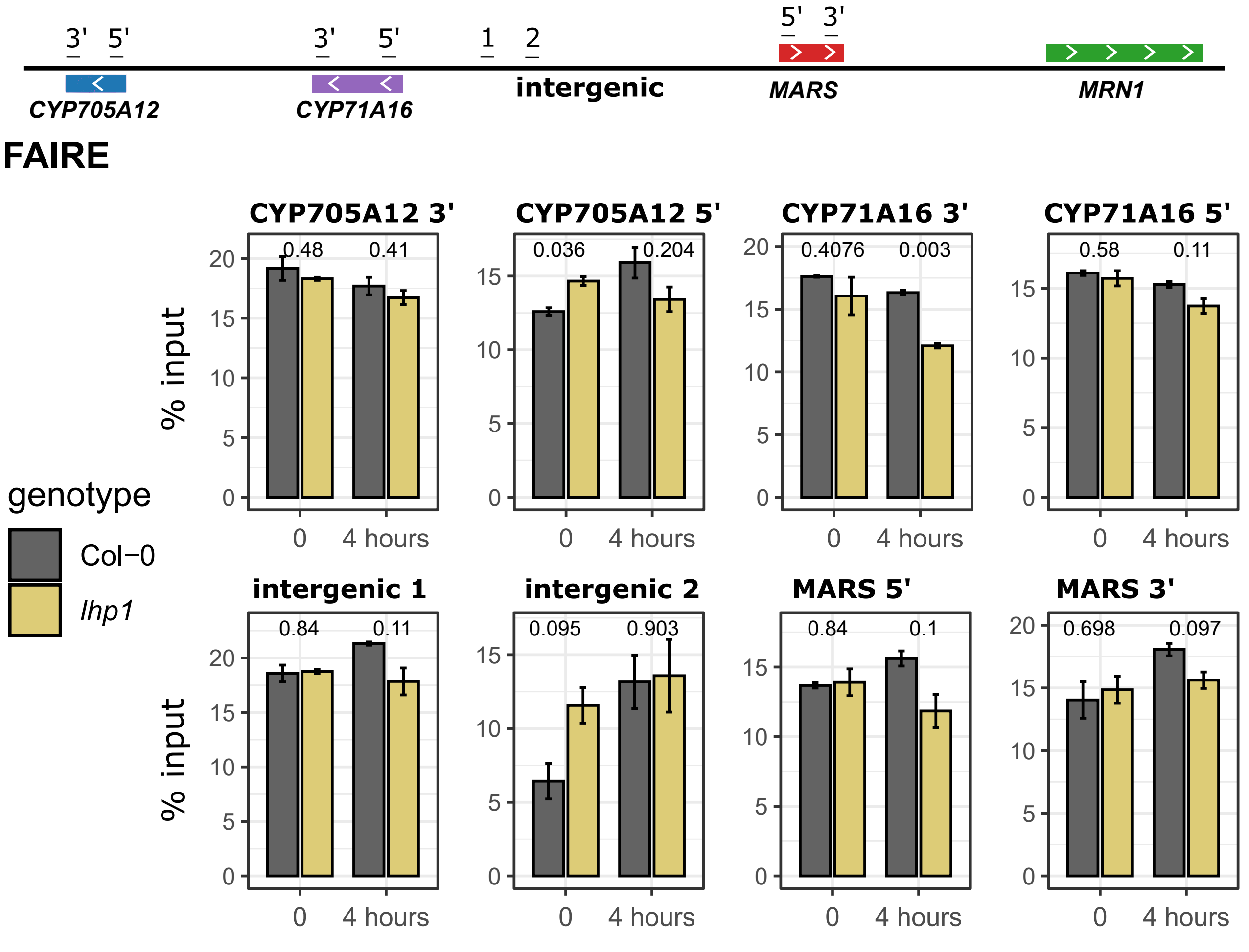

3'
5'
3'
5'
1
2
5'
3'
CYP705A12
CYP71A16
intergenic
MARS
MRN1
FAIRE
CYP705A12 3'
CYP705A12 5'
CYP71A16 3'
CYP71A16 5'
0.48
0.41
0.036
0.204
0.4076
0.003
0.58
0.11
intergenic 1
intergenic 2
MARS 5'
MARS 3'
0.84
0.11
0.095
0.903
0.84
0.1
0.698
0.097
% input
0
4 hours
genotype
Col−0
lhp1

### FigureS14

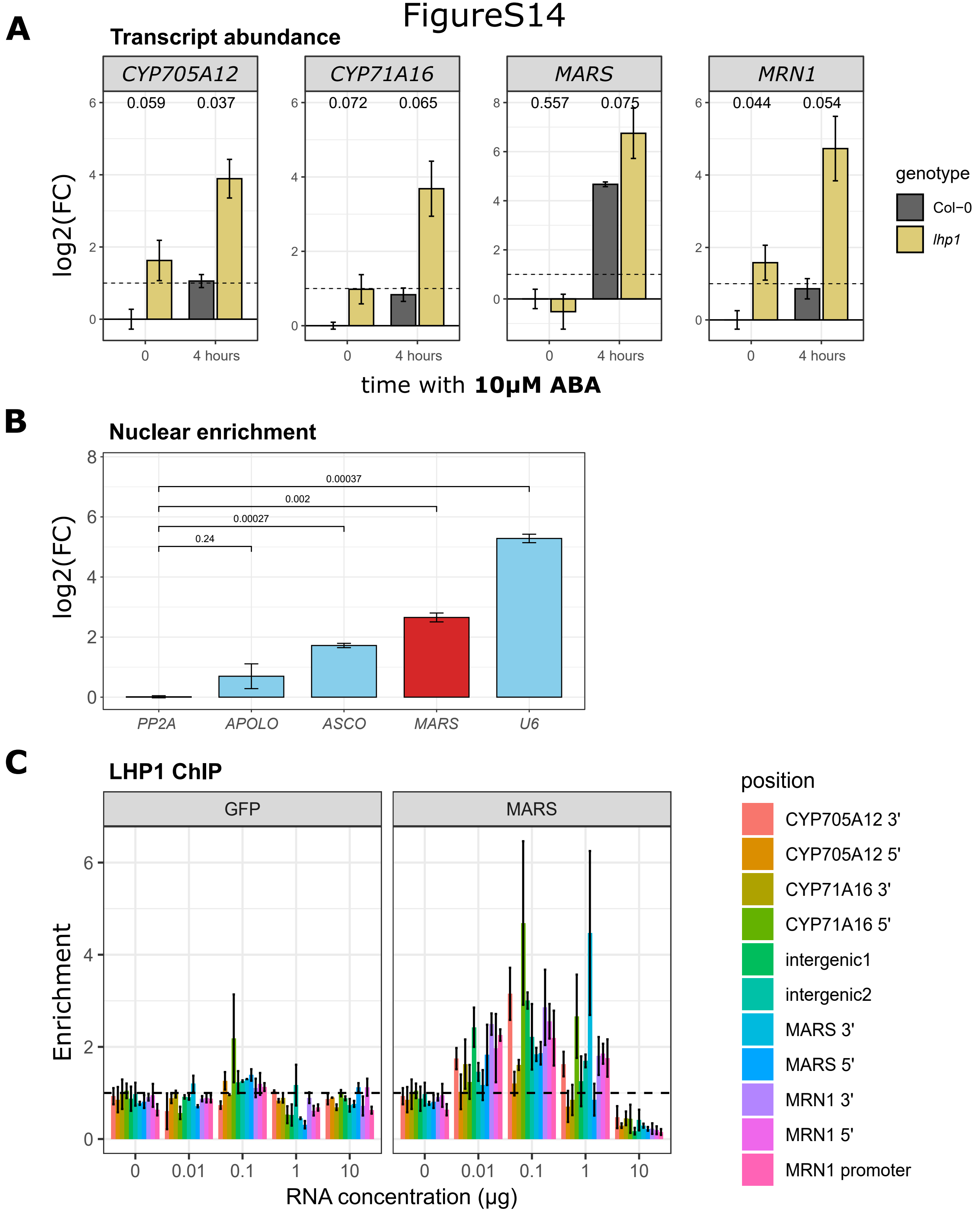
A
Transcript abundance
CYP705A12
CYP71A16
MARS
MRN1
0.059 0.037
0.072 0.065
0.557 0.075
0.044 0.054
log2(FC)
0
4 hours
time with 10μM ABA
genotype
Col−0
lhp1
B
Nuclear enrichment
log2(FC)
0.00037
0.002
0.00027
0.24
PP2A
APOLO
ASCO
MARS
U6
C
LHP1 ChIP
GFP
MARS
Enrichment
RNA concentration (μg)
position
CYP705A12 3'
CYP705A12 5'
CYP71A16 3'
CYP71A16 5'
intergenic1
intergenic2
MARS 3'
MARS 5'
MRN1 3'
MRN1 5'
MRN1 promoter

### FigureS15

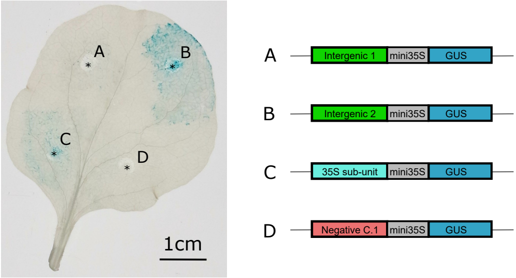
A
B
C
D
1cm
A
Intergenic 1
mini35S
GUS
B
Intergenic 2
mini35S
GUS
C
35S sub-unit
mini35S
GUS
D
Negative C.1
mini35S
GUS

| **Supplementary Table S1** | **List of primers used in the study** | |
|---|---|---|
| **Primers used to observe *MARS* isoforms** | ***Forward primer*** | ***Reverse primer*** |
| | CACCTCCTTCAGTGAGACCAGACGCT | TCAATGATAGGACTACTACTCATGGCCCAGG |
| **Primers used for *mrs1-1* (*SALK_133089)* homozygous selection** | ***Forward primer*** | ***Reverse primer*** |
| | AGTCCAGGTTTGGTTTGGTTC | ACATGTTCTTTTGGCAAGCAG |
| **Primers used to generate the RNAi lines** | ***Forward primer*** | ***Reverse primer*** |
| MARS | TCAATGATAGGACTACTACTCATGGCCCAGG | GATAGGACTACTACTCATGGCCCAGGTAAAC |
| **Primers used for transcripts abundance analysis from cDNA** | ***Forward primer*** | ***Reverse primer*** |
| *CYP705A12* | CAACAAGTGTTGTTTTCTCCGGTACG | GGGCTAAGATTGCATCCCTCGG |
| *CYP71A16* | CTGGTGTTTCTTTGTCGGCAGTTGT | CGGTCGGGGAATACATTCCGAGTT |
| *MARS* | ACTTTTTCACTGGTCGGTACCGAAGAC | TGCATAAAGTGTCGCACTCATTCGACTA |
| *MRN1* | GGGAGAAAGTGCTCTCTCTTGCCCTAA | GCGGCGCGATGAACAGGAGA |
| *AT1G13320 (housekeeping gene; Czechowski et al., 2005)* | GAGCTGAAGTGGCTTCCATGAC | GGTCCGACATACCCATGATCC |
| **Primers used for ChIP, meDIP and FAIRE experiments** | ***Forward primer*** | ***Reverse primer*** |
| CYP705A12.3' | GGAGTGCACTTAAGCGGATGAGCC | GCGATTGGAGCGATGGTGCAGT |
| CYP705A212.5' | GATGCGGAGATGGAGAAGAGGTCCA | CCTGCCTCCGAGCCCTCCTT |
| CYP71A16.3' | TGTGAGGTCGTTAAGCTCATGGAGAGG | GGAGCTGGAATCGATGTTTGCCGA |
| CYP71A16.5' | ATTGTGCCTTCCTCGCCGCT | TGGAGACGCTGGAAGAAGCAAGT |

| | | |
|---|---|---|
| intergenic1 | GGCTTTTGTTGACTACTATTACAGGCGGG | GGGTTTAGGGTTTATGGTTTAGGGTTTAGGG |
| intergenic2 | ACACCTATTAGTGACATCCACAAAGCGT | ACATTTTTGTGGTGTTAGGTGTGTGAAGC |
| MARS.5' | TTCCTTCAGTGAGACCAGACGCTTTCA | ACGGTTTCGCGTCCCGACC |
| MARS.3' | ACTTTTTCACTGGTCGGTACCGAAGAC | TGCATAAAGTGTCGCACTCATTCGACTA |
| MRN1.5' | TCGAGCAGAAGATCCCGAGAGTGAGA | CGTTAGCCTGACATGCCGCGT |
| MRN1.3' | GGGAGAAAGTGCTCTCTCTTGCCCTAA | GCGGCGCGATGAACAGGAGA |
| APOLO | GTGGCTTCCATAGCGCCGGA | CTAGCAACAGAGACCAACCC |
| **Primers used for *in-vitro* RNA transcription** | ***Forward primer*** | ***Reverse primer*** |
| *MARS* RNA | TAATACGACTCACTATAGGGGGCGACATTCACAAAACGTCGTTAAATA | GTTGGCAGCACCCATGTTTAGATGTCC |
| *GFP* RNA | GTAATACGACTCACTATAGGATGGTGAGCAAGGGCGAG | GGCTTGTACAGCTCGTCCATGC |
| **Primers used for transcript abundance analysis from sub-cellular localization and RIP experiments** | ***Forward primer*** | ***Reverse primer*** |
| *AT1G13320 (housekeeping gene; Czechowski et al., 2005)* | GAGCTGAAGTGGCTTCCATGAC | GGTCCGACATACCCATGATCC |
| APOLO | GTGGCTTCCATAGCGCCGGA | CTAGCAACAGAGACCAACCC |
| ASCO | CCCATCGCACTGATCGGCGG | TCGAGCGCTGCCGTCTTCAC |
| MARS | ACTTTTTCACTGGTCGGTACCGAAGAC | TGCATAAAGTGTCGCACTCATTCGACTA |
| U6 | CGGGGACATCCGATAAAATTGG | CGATTTGTGCGTGTCATCCTTG |
| MRN1 | GGGAGAAAGTGCTCTCTCTTGCCCTAA | GCGGCGCGATGAACAGGAGA |
| **Primers used for 3C experiment** | ***Forward primer*** | ***Reverse primer*** |
| Chromatin loop | GGCTTTTGTTGACTACTATTACAGGCGGG | CCAGACCAGTCATACACTCCTAGAACCTG |
| Control | GTCCGAATCTTACGGACCGGATTGTC | ACTGATAAACCCATCACCGGTGTTTCC |

| | | |
|---|---|---|
| **Primers used for the GUS transient assay** | ***Forward primer*** | ***Reverse primer*** |
| Negative control 1 | AACAGGTCTCAACCTAACTAAGTGTTACTAAATCATCTCACC | AACAGGTCTCTTGTTCATTTGTAAATCTTTTAAAGCCCTTG |
| Negative control 2 | AACAGGTCTCAACCTTGCTCAGAATGCCCCTACCT | AACAGGTCTCTTGTTCCTTGGTAACCCAAGCAACCA |
| Intergenic 1 | AACAGGTCTCAACCTGATTTTGGAGTTCCTGGTAAATGT | AACAGGTCTCTTGTTATGTCGCACTCATTCGACTA |
| Intergenic 2 | AACAGGTCTCAACCTGCATGTTGCCTATGATTAGAAGGA | AACAGGTCTCTTGTTACTCGATTTGGAGAAGTGTTTCAA |
| minimal 35S promoter element | AACAGGTCTCAAACAGCAAGACCCTTCCTCTATATAAGGAAGTTCATTTCATTTGGAGAGGACACGCTG | AACAGGTCTCTAGCCCAGCGTGTCCTCTCCAAATGAAATGAACTTCCTTATATAGAGGAAGGGTCTTGC |
| sub-unit B3 from 35S promoter element | AACAGGTCTCAACCTCATCGTTGAAGATGCCTCTGCCGACAGTGGTCCCAAAGATGGACCCCCACCCA | AACAGGTCTCTTGTTTGGGTGGGGGTCCATCTTTGGGACCACTGTCGGCAGAGGCATCTTCAACGATG |

#### LEGENDS TO SUPPLEMENTAL FIGURES

**Figure S1. *MARS* is coregulated with the genes of the marneral cluster**

Pearson correlation analysis derived from transcriptomics data from Araport11. Correlations between two genes are indicated with scores ranging from -1 to +1 where -1 corresponds to a negative correlation and +1 a positive correlation. A color scale indicates the Pearson correlation score. Each correlation was tested for significant differences (* for p ≤ 0.05, ** for p ≤ 0.01, *** for p ≤ 0.001).

**Figure S2. The RNAi-transgene does not affect DNA methylation at the *AT5G00580/MARS* locus.**

DNA methylation of the *AT5G00580/MARS* locus in Col-0 and RNAi-*MARS* seedlings assessed by MeDIP-qPCR under control condition. Higher values indicate 5mC enrichment. *APOLO* region has been taken as positive control of 5mC enrichment. Data are expressed as the mean ± standard error (n = 2) of the 5mC/IgG ratio. Letters indicate a statistical group determined by one-way analysis of variance (ANOVA) followed by Tukey's post-hoc test. For each genotype, letters indicate statistical difference between genomic position (p ≤ 0.05).

**Figure S3. *MARS* modulates the response of marneral genes to ABA without altering the plant sensitivity to an exogenous treatment**

(A) Average genotype effect on transcript levels of each marneral cluster gene in three independent RNAi-*MARS* lines compared to Col-0 in response to 10µM ABA according to two-way analysis of variance (ANOVA) including genotype and time as additive factors. Data are presented in Fig 2B. For each effect, numbers indicate the p-value of the difference between the RNAi lines and Col-0 by Tukey's post-hoc test.

(B) Transcript levels of two ABA marker genes, *RAB18* and *RD29B,* in response to 10µM ABA in RNAi-*MARS* lines. Gene expression data are expressed as the mean ± standard error (n = 3) of the log2 fold change compared to the Col-0 genotype at time 0h.

(C) Average genotype effect on the transcript levels of two ABA marker genes, *RAB18* and *RD29B,* in RNAi-*MARS* lines compared to Col-0 in response to 10µM ABA according to two-way analysis of variance (ANOVA) including genotype and time as additive factors. Data are presented in Fig S3B. For each effect, numbers indicate the p-value of the difference between the RNAi lines and Col-0 by Tukey's post-hoc test.

**Figure S4. *MARS* modulates the response of marneral genes to high concentrations of ABA**

(A) Transcript abundance of the genes of the marneral cluster in response to 100 µM ABA in RNAi-*MARS*. Gene expression data are expressed as the mean ± standard error (n = 3) of the log2 fold change compared to the Col-0 genotype at time 0h.

(B) Average genotype effect on the transcript levels of marneral cluster genes in RNAi-*MARS* compared to Col-0 in response to 100µM ABA according to two-way analysis of variance (ANOVA) including genotype and time as additive factors. Data are presented in Fig S4A. For each effect, numbers indicate the p-value of the difference between the RNAi lines and Col-0 by Tukey's post-hoc test.

(C) Transcript levels of two ABA marker genes in response to 100µM ABA in RNAi-*MARS* lines. Gene expression data are expressed as the mean ± standard error (n = 3) of the log2 fold change compared to the Col-0 genotype at time 0h.

(D) Average genotype effect on the transcript levels of two ABA marker genes in the different RNAi lines targeting AT5G00580/*MARS* compared to Col-0 in response to 100µM ABA according to two-way analysis of variance (ANOVA) including genotype and time as additive factors. Data are presented in

Fig S4C. For each effect, numbers indicate the p-value of the difference between the RNAi lines and Col-0 by Tukey's post-hoc test.

**Figure S5. Marneral cluster genes do not exhibit a circadian rhythm behavior**

(A) Transcript levels of the marneral cluster genes in response to water in RNAi-*MARS* lines. Gene expression data are expressed as the mean ± standard error (n = 3) of the log2 fold change compared to the Col-0 genotype at time 0h.

(B) Average genotype effect on the transcript levels of the marneral cluster genes in independent RNAi-*MARS* lines compared to Col-0 along water treatment according to two-way analysis of variance (ANOVA) including genotype and time as additive factors. Data are presented in Fig S5A. For each effect, numbers indicate the p-value of the difference between the RNAi lines and Col-0 by Tukey's post-hoc test.

(C) Transcript levels of two ABA marker genes, *RAB18* and *RD29B*, in response to water in RNAi-*MARS* lines. Gene expression data are expressed as the mean ± standard error (n = 3) of the log2 fold change compared to the Col-0 genotype at time 0h.

(D) Average genotype effect on the transcript levels of two ABA marker genes, *RAB18* and *RD29B*, in the independent RNAi-*MARS* lines compared to Col-0 in response to water treatment according to two-way analysis of variance (ANOVA) including genotype and time as additive factors. Data are presented in Fig S3A. For each effect, numbers indicate the p-value of the difference between the RNAi lines and Col-0 by Tukey's post-hoc test.

**Figure S6. Deregulation of *CYP71A16* does not modulate marneral cluster genes expression nor plant sensitivity to ABA**

(A) Transcript levels of the genes of the marneral cluster in response to 10µM ABA in *mro1-2 (CYP71A16* knock-out*)* mutant. Gene expression data are expressed as the mean ± standard error (n = 3) of the log2 fold change compared to the Col-0 genotype at time 0h.

(B) Average genotype effect on the transcript levels of the marneral cluster genes in the *mro1-2 (CYP71A16* knock-out*)* mutant compared to Col-0 in response to 10µM ABA according to two-way analysis of variance (ANOVA) including genotype and time as additive factors. Data are presented in Fig S6A. For each effect, numbers indicate the p-value of the difference between the *cyp71a16* mutant and Col-0 by Tukey's post-hoc test.

(C) Transcript levels of two ABA marker genes in response to 10µM ABA in *the mro1-2 (CYP71A16* knock-out*)* mutant. Gene expression data are expressed as the mean ± standard error (n = 3) of the log2 fold change compared to the Col-0 genotype at time 0h.

(D) Average genotype effect on the transcript levels of two ABA marker genes in the *mro1-2 (CYP71A16* knock-out*)* mutant compared to Col-0 in response to 10µM ABA according to two-way analysis of variance (ANOVA) including genotype and time as additive factors. Data are presented in Fig S6C. For each effect, numbers indicate the p-value of the difference between the *cyp71a16* mutant and Col-0 by Tukey's post-hoc test.

**Figure S7. Deregulation of *MRN1* does not modulate marneral cluster genes expression nor plant sensitivity to ABA**

(A) Transcript levels of the genes of the marneral cluster in response to 10µM ABA in *35S:MRN1* and *mrn1* mutants. Gene expression data are expressed as the mean ± standard error (n = 3) of the log2 fold change compared to the Col-0 genotype at time 0h.

(B) Average genotype effect on the transcript levels of the genes of the marneral cluster in *35S:MRN1* and *mrn1* mutants compared to Col-0 in response to 10µM ABA according to two-way analysis of variance (ANOVA) including genotype and time as additive factors. Data are presented in Fig S7A. For each effect, numbers indicate the p-value of the difference between the *MRN1* mutants and Col-0 by Tukey's post-hoc test.

(C) Transcript levels of the genes of two ABA marker genes, *RAB18* and *RD29B*, in response to 10µM ABA in *35S:MRN1* and *mrn1* mutants. Gene expression data are expressed as the mean ± standard error (n = 3) of the log2 fold change compared to the Col-0 genotype at time 0h.

(D) Average genotype effect on the transcript levels of two ABA marker genes, *RAB18* and *RD29B*, in *35S:MRN1* and *mrn1* mutants compared to Col-0 in response to 10µM ABA according to two-way analysis of variance (ANOVA) including genotype and time as additive factors. Data are presented in Fig S7C. For each effect, numbers indicate the p-value of the difference between the *MRN1* mutants and Col-0 by Tukey's post-hoc test.

**Figure S8. *MARS* modulates seed germination and mannitol-dependent root growth through the regulation of *MRN1* expression**

(A) Percentage of germinated seeds in a control medium. Results are expressed as the mean ±standard error (n = 7) from a batch of ≈50 seeds collected from plants grown separately. Time for 50% germination (T50) is indicated on the right. Letters indicate a statistical group determined by one-way analysis of variance (ANOVA) followed by Tukey's post-hoc test. For each genotype, letters indicate statistical difference between T50 ($p \leq 0.05$).

(B) Percentage of germinated seeds in a medium containing 0.5µM ABA. Results are expressed as the mean ±standard error (n = 7) from a batch of ≈50 seeds collected from plants grown separately. Time for 50% germination (T50) is indicated on the right. Letters indicate a statistical group determined by one-way analysis of variance (ANOVA) followed by Tukey's post-hoc test. For each genotype, letters indicate statistical difference between T50 ($p \leq 0.05$).

(C) Percentage of germinated seeds in a control medium. Results are expressed as the mean ±standard error (n = 4) from a batch of ≈50 seeds collected from plants grown separately. Time for 50% germination (T50) is indicated on the right. Letters indicate a statistical group determined by one-way analysis of variance (ANOVA) followed by Tukey's post-hoc test. For each genotype, letters indicate statistical difference between T50 ($p \leq 0.05$).

(D) Percentage of germinated seeds in a medium containing 0.5µM ABA. Results are expressed as the mean ±standard error (n = 4) from a batch of ≈50 seeds collected from plants grown separately. Time for 50% germination (T50) is indicated on the right. Letters indicate statistic group determined by one-way analysis of variance (ANOVA) followed by Tukey's post-hoc test. For each genotype, letters indicate statistical difference between T50 ($p \leq 0.05$).

(E) Mean primary root length, lateral root length and lateral root density according to the genotype and the condition of 11-day-old seedlings. Letters indicate a statistical group determined by one-way analysis of variance (ANOVA) followed by Tukey's post-hoc test. For each condition, letters indicate statistical difference between genotypes ($p \leq 0.05$).

**Figure S9. The epigenetic landscape of the marneral cluster and surrounding genomic region**

First track represents DNA accessibility determined by the ATAC-seq (44). ATAC-peaks are indicated with a blue rectangle and correspond to relaxed chromatin. Second and third tracks show H3K27me3 and LHP1 ChIP-Seq, respectively (43). The three experiments shown here have been performed using Arabidopsis shoot. Gene annotation is shown at the top.

**Figure S10. *MARS* influences H3K27me3 deposition in the marneral cluster region**

H3K27me3 deposition in the intergenic region, *CYPs* and *MARS* loci was measured in Col-0 and RNAi-*MARS* seedlings was assessed by ChIP-qPCR in control conditions and in response to ABA. Higher values indicate H3K27me3 enrichment. Values under the dotted line are considered as not enriched.

Data are expressed as the mean ± standard error (n = 2) of the H3K27me3/Igg ratio. Numbers are p-value of the difference between the two genotypes determined by Student t-test.

**Figure S11. *MARS* modulates LHP1 binding across the marneral cluster region**

LHP1 binding to the intergenic region, *CYPs* and *MARS* loci was measured in Col-0 and RNAi-*MARS* seedlings was assessed by ChIP-qPCR in control conditions and in response to ABA. Higher values indicate LHP1 enrichment. Values under the dotted line are considered as not enriched. Data are expressed as the mean ± standard error (n = 2) of the LHP1/Igg ratio. Numbers are p-value of the difference between the two genotypes determined by Student t-test.

**Figure S12. *MARS* modulates chromatin condensation of the marneral cluster genomic region**

Chromatin condensation in the intergenic region, *CYPs* and *MARS* loci was measured in Col-0 and RNAi-*MARS* seedlings was assessed by Formaldehyde Assisted Isolation of Regulatory Element (FAIRE)-qPCR in control conditions and in response to ABA. Lower value indicates more condensed chromatin. Results are expressed as the mean ± standard error (n = 2) of the percentage of input (signal measured before isolation of decondensed region of chromatin; free of nucleosomes). Numbers are p-value of the difference between the two genotypes determined by Student t-test.

**Figure S13. LHP1 is involved in chromatin condensation modulation of the marneral cluster region**

Chromatin condensation in the intergenic region, *CYPs* and *MARS* loci was measured in Col-0 and *lhp1* mutant seedlings was assessed by Formaldehyde Assisted Isolation of Regulatory Element (FAIRE)-qPCR in control conditions and in response to ABA. Lower value indicates more condensed chromatin. Results are expressed as the mean ± standard error (n = 3) of the percentage of input (signal measured before isolation of decondensed region of chromatin; free of nucleosomes). Numbers are p-value of the difference between the two genotypes determined by Student t-test.

**Figure S14. Nuclear-enriched *MARS* RNA modulates LHP1 binding to the marneral cluster and influences the subsequent ABA response of the marneral cluster genes**

(A) Transcript levels of the marneral cluster genes in response to ABA treatment in *lhp1* mutant. Results are expressed as the mean ± standard error (n = 3) of the log2 fold change compared to time-point 0h. Numbers are p-value of the difference between the two genotypes determined by Student t-test.

(B) Nuclear enrichment of the lncRNA *MARS* compared to other nuclear-enriched lncRNAs determined as the ratio of transcript abundance in the nuclear fraction compared to total cellular RNA. Higher value indicates nuclear enrichment. *APOLO*, *ASCO* and *U6* RNA have been used as positive controls whereas *RHIP1* (*AT4G2641;* housekeeping gene) has been used as negative control. Results are expressed as the mean ± standard error (n = 3) of the log2 fold change compared to the total cell fraction. Numbers are p-value of the difference between the corresponding RNA determined by Student t-test.

(C) LHP1 binding in RNAi-*MARS*-derived chromatin to different sites across the marneral cluster upon increasing amounts of *in-vitro* transcribed *MARS* or GFP RNA (ug), determined by ChIP-qPCR. Higher values indicate LHP1 enrichment. Results are expressed as the mean ± standard error (n = 2) of the LHP1/Igg ratio.

**Figure S15. The intergenic region between *CYP71A16* and *MARS* is able to activate gene transcription**

Agroinfiltration of the different constructs from Figure 5D in the same tobacco leaf. Points of agroinfiltration for each construct are indicated with a star (*), and each letter indicates the corresponding constructs. Same result has been observed in two other independent tobacco leaves.
